# Phosphorylation of the Smooth Muscle Master Splicing Regulator RBPMS Regulates its Splicing Activity

**DOI:** 10.1101/2022.08.11.503562

**Authors:** Michael D. Barnhart, Yi Yang, Erick E. Nakagaki-Silva, Thomas H. Hammond, Mariavittoria Pizzinga, Clare Gooding, Katherine Stott, Christopher W.J. Smith

## Abstract

We previously identified RBPMS as a master regulator of alternative splicing in differentiated smooth muscle cells (SMCs). *RBPMS* is transcriptionally downregulated during SMC dedifferentiation, but we hypothesized that RBPMS protein activity might be acutely downregulated by post-translational modifications. Publicly available phosphoproteomic datasets reveal that Thr113 and Thr118 immediately adjacent to the RRM domain are commonly both phosphorylated. An RBPMS T113/118 phosphomimetic T/E mutant showed decreased splicing regulatory activity both in transfected cells and in a cell-free *in vitro* assay, while a non-phosphorylatable T/A mutant retained full activity. Loss of splicing activity was associated with a modest reduction in RNA affinity but significantly reduced RNA binding in nuclear extract. A lower degree of oligomerization of the T/E mutant might cause lower avidity of multivalent RNA binding. However, NMR analysis also revealed that the T113/118E peptide acts as an RNA mimic which can loop back and antagonize RNA-binding by the RRM domain. Finally, we identified ERK2 as the most likely kinase responsible for phosphorylation at Thr113 and Thr118. Collectively, our data identify a potential mechanism for rapid modulation of the SMC splicing program in response to external signals during the vascular injury response and atherogenesis.

## INTRODUCTION

Noncommunicable diseases (NCDs), specifically cardiovascular diseases (CVDs), are the leading cause of global death rates. Of the 56.9 million global deaths in 2016, 40.5 million were due to NCDs, of which the leading cause of NCDs was CVD totaling 17.9 million, with more than 75% of these deaths occurring in low- and middle-income countries (WHO, 2019). Pathological blood vessel narrowing carries the highest mortality rate of all the CVDs as a result of the dysregulation of vascular smooth muscle cells (VSMCs), a response that cannot yet be therapeutically controlled. Thus, novel therapeutics and early diagnostic methods at low costs are imperative to reduce the economic burden these diseases carry (3) and it is vital to develop a deeper understanding of CVD mechanisms, particularly the molecular mechanisms underlying the control of VSMCs.

Dysfunctional VSMCs drive several CVDs such as atherosclerosis and restenosis, cerebral microangiopathy, Marfan syndrome, hypertension, and pulmonary arterial hypertension (3). Coronary artery diseases (CADs), atherosclerosis and restenosis, are the most common CVDs and thus the major cause of death worldwide. During atherosclerosis and restenosis, VSMCs are stimulated to proliferate by platelet-derived growth factor (PDGF) driving the infiltration and migration of VSMCs to the intima and resulting in the progressive thickening and narrowing of the blood vessel lumen (4), ultimately leading to ischemic heart disease and stroke. In fact, VSMC phenotypic dysregulation is the underlying cause of each of the aforementioned diseases. Therefore, understanding this VSMC phenotypic switch from quiescent-contractile to migratory-proliferative on the molecular level is critical and must be pursued. While the transcriptional program underpinning VSMC phenotypic switching has been well studied, the accompanying program of regulated alternative splicing (AS), and the RNA-binding proteins (RBPs) responsible for regulating it, are less well characterized.

To this end, RBPMS (<u>R</u>NA <u>B</u>inding <u>P</u>rotein with <u>M</u>ultiple <u>S</u>plicing) was recently identified as a potential SMC master splicing regulator, responsible for controlling numerous components critical to differentiated VSMC function (5). RBPMS was found to be highly expressed in the differentiated phenotype and not detected in the dedifferentiated (proliferative) phenotype, suggesting a regulatory role in differentiated SMCs (5). In differentiated SMCs, RBPMS was found to both repress exons included within the dedifferentiated phenotype by binding upstream or within the repressed exons (found in transcripts such as *ACTN1* and *TPM1*) and activate the inclusion of exons found in the differentiated phenotype by binding downstream of the included exon (found in transcripts such as *FLNB, MPRIP*, and *PTPRF*) (5). RBPMS has a single RRM domain that mediates both RNA binding to CAC motifs as well as dimerization followed by an intrinsically disordered C-terminal tail of 84 amino acids in the canonical RBPMS-A isoform (Figure 1A). The reported binding motif of RBPMS consists of tandem CACs with a variable spacer of approximately 1-12 nucleotides, consistent with the dimeric structure (6, 7). The canonical isoform, RBPMS-A, was found to be more active in overall splicing regulation compared to the other isoforms; in particular, RBPMS-A could both repress and activate target exons while RBPMS-B was only able to activate.

Upon knockdown of RBPMS in differentiated PAC1 cells (a well-established SM cell line), alterations of alternative splicing in numerous genes encoding components of the actin cytoskeleton and cell adhesion machinery were accompanied by reorganization of actin structures to resemble the dedifferentiated phenotype (5). Moreover, overexpression of RBPMS in PAC1 cells was able to recapitulate some differentiated SMC AS patterns that are usually only observed in fully differentiated tissue SMCs, but not in cell culture. Moreover, the SMC transcription factor Myocardin was recently shown to drive the SMC AS program in part by up-regulating RBPMS transcription (8). These findings demonstrate that RBPMS is a master regulator of the alternative splicing program found in differentiated smooth muscle cells. Thus, the maintenance of the differentiated state is controlled by both transcriptional regulation and alternative splicing regulation (5). Therefore, the precise control of this splicing regulator is vital in determining SMC fate. RBPMS was identified as a candidate master regulator based on the super-enhancer associated with the *RBPMS* gene in SMC-rich tissues, and it is transcriptionally downregulated in phenotypic modulation (5). However, transcriptional down-regulation does not provide for rapid downregulation of the activity of proteins already present in the cell.

Various growth factors activate the dedifferentiation of VSMCs by initiating the classical Ras/Raf/MEK/ERK pathway which leads to the downstream phosphorylation of several different families of proteins (3). Phosphorylation of RNA-binding proteins (RBPs), specifically splicing regulators, has been shown to affect proteins in different ways including, but not limited to, protein function, localization, and degradation (9–12). We were thus interested in the possibility that upon the onset of phenotypic transition, phosphorylation might play a role in the acute regulation of RBPMS protein activity. Therefore, we set out to determine if RBPMS could be phosphorylated and, if so, what role this plays in modulating its activity. We found that ERK2 binds to an F-site domain located on the C-terminal region of RBPMS and phosphorylates RBPMS-A at Thr113, Thr118, just downstream of the RRM domain (Figure 1A), and Ser144. The use of phosphomimetic mutants indicated that phosphorylation of Thr113 and Thr118, but not Ser144, results in the inhibition of RBPMS-A splicing activity which leads to a splicing profile found in proliferative SMCs. Furthermore, phosphorylation of Thr113 and Thr118 leads to reduced RNA-binding due to both a reduction in oligomerization and associated avidity of multivalent RNA interaction as well as a direct occlusion of the RNA-binding surface of the RRM domain, as elucidated via biochemical assays and NMR. Overall, our data provide a molecular basis for the rapid modulation of RBPMS-A activity by phosphorylation in response to external signals during the vascular injury response and atherogenesis.

**Figure 1.**
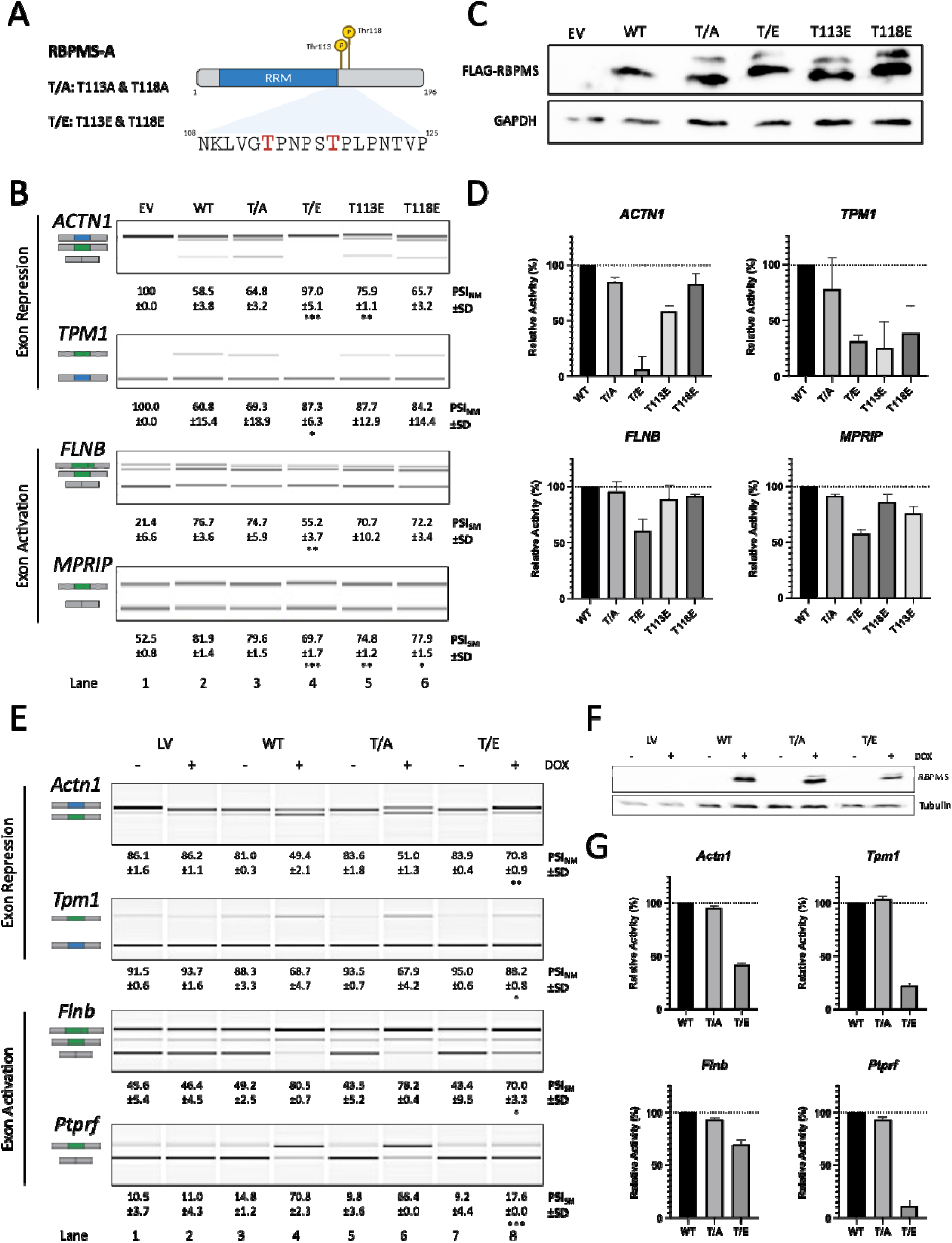
RBPMS-A Phosphomimetics Have Reduced Splicing Activity Across Cell Lines. (A) Graphical representation of RBPMS-A with the T113 and T118 highlighted in red. T/A is a double mutant of T113A and T118A, while T/E is double mutant of T113E and T118E. Created with BioRender.com (B) Splicing analysis of the exon repression function of wild-type (WT) RBPMS-A and effectors in endogenous *ACTN1* and *Tpm1*, and its exon activation function in endogenous *FLNB* and *MPRIP* in HEK293T cells. RT-PCR followed by visualization on QIAxcel. Cartoons below the gene names represent the PCR products. The blue exon in the cartoon represents the NM (Non-smooth Muscle) exon, while the green exon represents the SM (Smooth Muscle) exon. PSI (Percent Spliced In) values are the mean ± SD (n = 3). A Student’s t-test was used to determine statistical significance between wild-type RBPMS-A and effector. EV=empty vector (C) Immunoblot of FLAG-RBPMS effectors used for splicing assays in (B). Whole-cell lysates were probed with antibodies to FLAG and GAPDH. GAPDH was used as a loading control. (D) Bar graphs showing the percent relative splicing efficiency of each effector normalized to wild-type RBPMS-A (black bar) after background splicing was taken into consideration. (E) Splicing analysis of the exon repression function of wild-type (WT) RBPMS-A and effectors in endogenous *Actn1* and *Tpm1*, and its exon activation function in endogenous *Flnb* and *Ptprf* in rat PAC1 cells. RT-PCR followed by visualization on QIAxcel. Cartoons below the gene names represent the PCR products. The blue exon in the cartoon represents the NM (Non-smooth Muscle) exon, while the green exon represents the SM (Smooth Muscle) exon. PSI (Percent Spliced In) values are the mean ± SD (n = 3). A Student’s t-test was used to determine statistical significance between wild-type RBPMS-A and effector. LV = empty lentiviral vector. (F) Immunoblot of Dox inducible RBPMS effectors used for splicing assays in (E). Whole-cell lysates were probed with antibodies to RBPMS and Tubulin. Tubulin was used as a loading control. (G) Bar graphs showing the percent relative splicing efficiency of each effector normalized to wild-type RBPMS-A (black bar), after background splicing was taken into consideration. Error bars indicate SD. Significance in panels B and E indicated by *p < 0.05, **p < 0.01, ***p < 0.001.

## MATERIALS AND METHODS

### Growth conditions for bacterial cultures

*Escherichia coli (E. coli)* grown on lysogeny broth (LB) agar was incubated overnight at 37°C, while *E. coli* grown in LB liquid media was incubated at 37°C overnight (or for a minimum of eight hours) and agitated at 280 RPM. Both media contained the appropriate antibiotic; the final concentration of ampicillin used was 50 μg/mL, while kanamycin was 30 μg/mL. For small-scale (miniprep) DNA purification, 5 mL of LB was used, while large-scale (maxiprep) required 100 mL.

### Transformation of *E. coli*

Laboratory stocks of chemically competent *E. coli* cells (TG-1) stored at -80°C were used for bacterial transformation via heat shock. 100 μL of TG-1 cells were added and mixed with 100 ng of plasmid DNA or 2.5 μL of a ligation reaction (see below) and incubated on ice for 30 minutes. The mix was then heat shocked at 42°C for 30 seconds and then placed back on ice for five minutes. Finally, the entire mix was spread on LB agar plates containing the appropriate antibiotic and incubated as described above.

### Ligation of DNA fragments

Ligation reactions were set up in a 1:3 ratio of vector:insert, calculated from base-pair length. T4 DNA ligase (Roche) was used, and the final reaction volume was 10 μL. Ligation reactions were then used for *E. coli* transformations as described above.

### Plasmid DNA purification

Following transformation, single colonies of *E. coli* were picked from the LB agar plates and placed in either 5 mL LB for miniprep or 100 mL for maxiprep with the appropriate antibiotic. Cultures were incubated at 37°C, agitating at 280 RMP, for at least 8 hours for minipreps and overnight for maxipreps. Bacterial cells were then pelleted at either 4800 × g for 5 mins for minipreps and 5000 × g for 15 mins for maxipreps at 4°C. Manufacturer’s instructions for the QIAGEN QIAprep Spin Miniprep or Maxiprep kits were then followed for cell lysis and plasmid DNA purification. A NanoDrop spectrometer (Thermo Scientific) was used to measure the final purified plasmid DNA concentration and appropriate dilutions were made with ddH_2_O and stored at -20°C.

### Purification of DNA fragments

DNA fragments that were amplified by PCR were purified using a QIAGEN QIAquick PCR Purification Kit. If restriction digestion was necessary, the DNA fragment was run on an agarose gel and then excised and purified using a QIAGEN QIAquick Gel Extraction Kit. DNA fragments were eluted with ddH_2_O and stored at -20°C.

### DNA construct generation

RBPMS-A constructs (wild-type and some mutant) were previously generated in the lab and cloned into pEGFP-C1. However, to reduce the size of the N-terminal tag, all constructs were cloned into either pCI-3xFLAG at *EcoRI* and *BamHI* or pCI-3xFLAG-NLS at *Xho1* and *BamHI* for use *in vivo* overexpression in HEK293T cells. An exogenous nuclear localization signal (NLS) was used to ensure the nuclear localization of the deletion mutants, given that RBPMS NLS is not well characterized.

Oligonucleotides for the desired mutagenesis of RBPMS were designed using the Agilent QuikChange Primer Design software available online followed by mutagenesis PCR according to the manufacturer’s guidelines. Deletion mutants were generated by designing oligonucleotide primers that amplified in the opposite direction, leaving the desired deletion omitted from amplification. The blunt ends that were generated were then phosphorylated and ligated. Parental plasmids in both mutant and deletion generation were digested using *DpnI* enzyme.

RBPMS-A and various mutants were subcloned into a modified pET15b-TEV vector (Yi Yang et al., manuscript in preparation) at *SalI* and *XhoI* for the expression of recombinant protein in *E. coli*. The pET15b vector was modified by replacing the thrombin site with a TEV site and replacing the *NdeI* site with *SlaI* because both sites are present in RBPMS-A.

MEK1, CA-MEK1, and ERK2 were subcloned from pIVEX_MEK1 (WT), pET28a_MEK1_del44-51_S218D_M219D_N221D_S222D (Highly Constitutively Active), and pET28a_ERK2 (WT), respectively, and cloned into pCI-3xFLAG at *AvrII* and *XhoI*. These constructs were obtained from Remkes Scheele, Department of Biochemistry, University of Cambridge.

The sequences of each construct were verified by Sanger sequencing. The oligonucleotides used to create each construct can be found in Table 1.

**Table 1.**
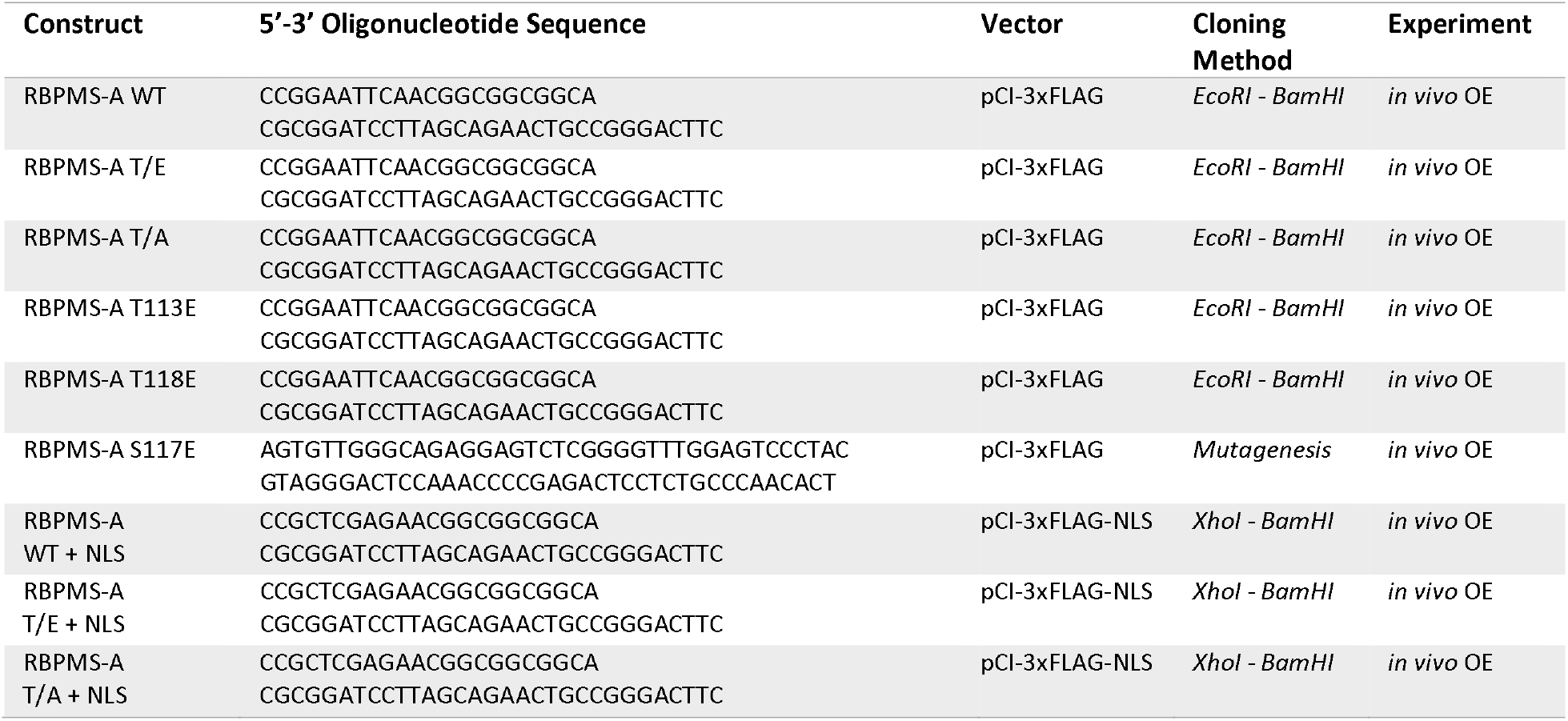
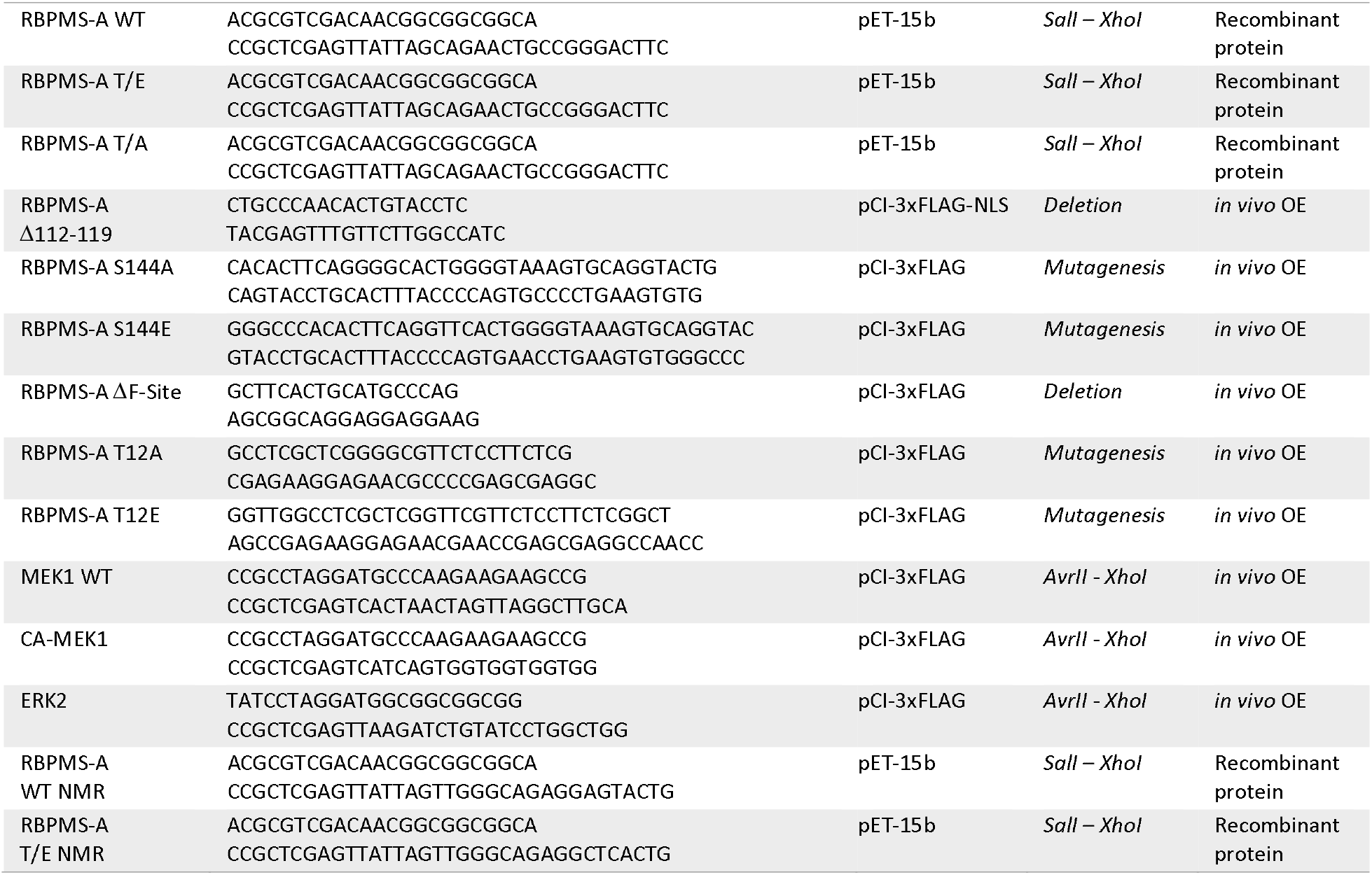
. Oligonucleotide sequences used for the generation of DNA constructs

### Cell Culture

The rat Pulmonary Artery Cell (PAC1) line was used in this study because they are a well-established smooth muscle cell model (Rothman et al., 1992). Additionally, the human embryonic kidney cell line with a mutated version of the SV40 large T antigen (HEK 293T) was used as an alternative cell line due to the poor transfection efficiency of PAC1 cells.

Both cell lines were cultured in Dulbecco’s Modified Eagle Medium (DMEM, Gibco) containing 1% Glutamax and 10% fetal bovine serum (FBS). Cells were cultured under standard conditions, 37°C and 5% CO_2_. HEK293T cells were passaged twice a week at a 1:15 dilution, while PAC1 cells were only passaged once a week at a 1:10 dilution to maintain the differentiated state and avoid stimulating them to become more proliferative.

### Transient Transfections

HEK293T cells were seeded in a six-well plate at a concentration of 2x10^5^ cells/well at least 8 hours prior to transfection. For each well, 2 μL of Lipofectamine 2000 reagent (Thermo Fisher) and 1 μg of DNA were diluted in Opti-MEM (Gibco). Cells were incubated under standard conditions (described above) for 48 hours at which time total RNA and protein were harvested.

### Inducible lentiviral overexpression

The inducible lentiviral overexpression cell lines used in this study were created using pInducer22 plasmids and the Gateway System (13). Overexpression was achieved by inducing the cells with 1 μg/mL of doxycycline diluted in the cell culture medium. Total RNA and protein were harvested 24 hours post-induction.

### Western Blotting

Western blotting was used to verify the protein expression levels of the overexpression experiments. Total lysates were harvested by the direct addition of 1xSDS + 5% β-mercaptoethanol and stored at -80°C. Lysates were then boiled at 95°C, run on an SDS-PAGE gel, and transferred to an Immobilon-FL membrane (Millipore). Specific proteins were detected by the primary antibodies listed in Table 2. Donkey anti-mouse IgG antibodies (Stratech Scientifica) conjugated to horseradish peroxidase were used at a 1:10,000 dilution. Antibody detection was carried out using luminol/iodophenol chemiluminescence followed by imaging on a Bio-Rad ChemiDoc and analysis on Image Lab (v6.0).

**Table 2.** . Antibodies used in this study for Western Blotting

| **Protein** | **Antibody** | **Concentration** |
| --- | --- | --- |
| FLAG | Sigma Aldrich, F-1804 | 1:2000 |
| GAPDH | Santa Cruz, sc-47724 | 1:1000 |

### Immunofluorescence

HEK293T cells were seeded on a Corning BioCoat Poly-D-Lysine 8-well culture slide at a concentration of 14x10^3^ cells/well at least 8 hours prior to transfection. 48 hours post-transfection, slides were incubated in 3.8% paraformaldehyde (PFA) for five minutes then rinsed with phosphate-buffered saline (PBS). Cells were then permeabilized with 0.5% NP-40 for two minutes and rinsed with PBS. 1% bovine serum albumin (BSA) in PBS (blocking buffer) was used to block the cells for one hour and replaced with anti-FLAG antibodies (Sigma Aldrich, F-1804) diluted in blocking buffer (1:1000) for an additional hour. Next, cells were rinsed with PBS and incubated with anti-mouse Rhodamine Red-X (Thermo Fisher, R-6393) diluted in blocking buffer (1:500) for one hour. Following a final rinse with PBS, the chambers were detached from the slides, and coverslips were mounted using ProLong Diamond Antifade with DAPI (Thermo Fisher). All incubations were carried out at room temperature. Images were acquired on a Zeiss Axio Observer.Z1 LSM 980 microscope equipped with Airyscan 2, using ZEN Blue software (version 3.3), in Airyscan super-resolution imaging mode. A C-Apochromat 40X /1.20 W Korr objective was used. 15/20 Z-slices per image were acquired at an interval of 0.2 µm. Airyscan processing with standard parameters was applied to each image in Zen Blue software. Further processing and analysis were performed using FiJi software (version 2.3.051). Supplementary data images of cells were visualized and acquired using an EVOS M5000 with a 40X objective and analyzed on EVOS M5000 software (v1.3).

### RT-PCR

TRI reagent (Sigma) was used to extract total RNA from cells following the manufacturer’s guidelines. Genomic DNA was then degraded using Turbo DNase (Thermo Fisher) and RNA was quantified on a NanoDrop spectrometer (Thermo Fisher). 1 μg of total RNA along with oligo dTs and SuperScriptII (Thermo Fisher) were used to prepare cDNA following the manufacturer’s protocol.

### Alternative splicing analysis

To detect changes in splicing isoforms, PCR reactions were prepared with 1 μL of cDNAs (described above), oligonucleotides targeting specific genes (Table 3), and Jumpstart Taq DNA polymerase (Sigma). The following PCR parameters were used: initial denaturation at 94°C for 3 min; 35 cycles of 94°C for 30 sec, 60°C for 30 sec, and 72°C for 60 sec; final elongation at 94°C for 30 secs. Reaction was then slow cooled, ΔT -1°C every min down to 37°C.

**Table 3.**
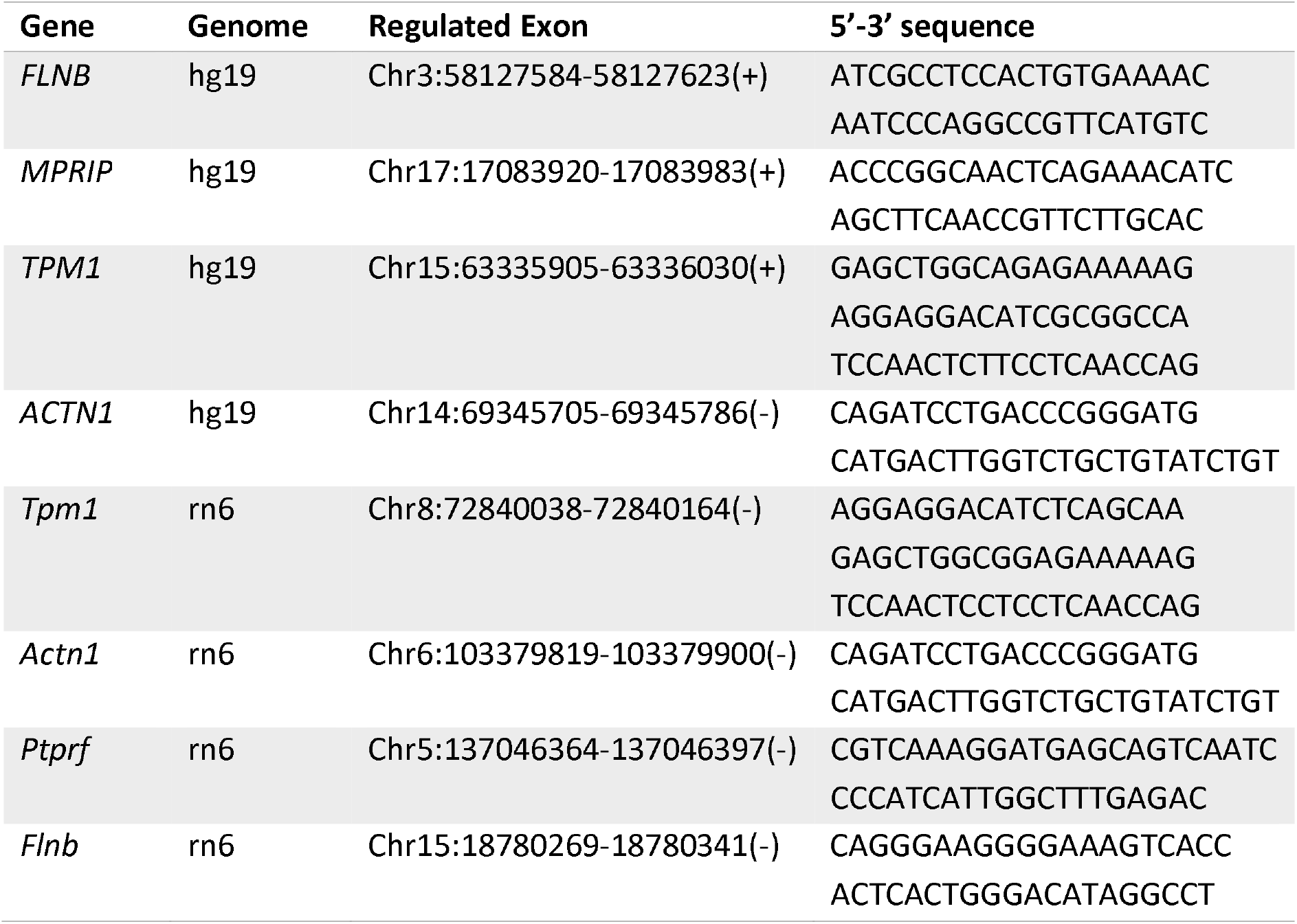
. Oligonucleotide sequences used to study alternative splicing

QIAxcel Advanced System (QIAGEN) was used to separate the PCR products and QIAxcel ScreenGel software was used to analyze the PCR products and calculate the percentage spliced in (PSI) value. PSIs are shown as the mean (%) ± standard deviation of at least three experiments. Negative controls included a no template (cDNA) reaction and minus RT reaction.

### Recombinant Protein Expression

RBPMS-A wild-type and mutants were cloned into modified pET15b vectors containing an N-terminal His_6_ tag and a TEV protease site. BL21 competent cells were then transformed with each construct and a single colony was then inoculated in 10 mL of LB + ampicillin (50 μg/mL) and grown for 12 hours at 37°C, 280 RPM. This cell slurry was then added to 500 mL of LB + ampicillin (50 μg/mL) and incubated at 37°C, 220 RPM for approximately 2 hours until the OD reached 0.6-0.8 (log phase growth). 0.2 mM of IPTG (isopropyl β-D-1-thiogalactopyranoside) was added to induce the expression of protein for 2 hours at 37°C, 220 RPM. Cells were then pelleted at 7,000 x g for 10 minutes at 4°C and resuspended in 10 mL of Buffer A (50 mM Tris Base, 0.5 M KCl, 40 mM imidazole, 10% glycerol, pH 8.5 at 4°C). 1 mM DTT, 1.6 M LiCl, and 2 cOmplete^TM^ mini protease inhibitor cocktail (Roche) tablets were added to the cell suspension followed by cell lysis using a French Press (Stansted). Lysates were centrifuged at 40,000 x g for 30 minutes at 4°C for the separation of soluble and insoluble fractions. Soluble fractions were filtered using a 0.45 μM filter in preparation of protein purification.

### Purification of RBPMS

Filtered soluble recombinant wild-type RBPMS-A and mutants were purified on an AKTA protein purification system (GE Healthcare;). Recombinant protein was first purified on HisTrap HP column (GE Healthcare) using Buffer A (50mM Tris Base, 0.5 M KCl, 40 mM imidazole, 10% glycerol, 1 mM DTT, pH 8.5 at 4°C), Buffer A2 (50 mM Tris Base, 2 M NaCl, 10% glycerol, 1 mM DTT, pH 8.5 at 4°C), and Buffer B (50mM Tris Base, 0.5 M KCl, 500 mM imidazole, 10% glycerol, 1 mM DTT, pH 8.5 at 4°C). After fraction analysis on SDS-PAGE, the fractions containing recombinant protein were buffer exchanged to Buffer QA (25 mM CAPS, 50 mM KCl, 10% glycerol, pH 10 at 4°C) followed by TEV protease cleavage at a ratio of 1:40 for 2 hours at 30°C and stored overnight at 4°C. Cleaved recombinant protein was then purified on a MonoQ 5/50 GL column (GE Healthcare) using Buffer QA (described above + 1 mM DTT) and Buffer QB (25 mM CAPS, 50 mM KCl, 1 M NaCl, 10% glycerol, 1 mM DTT, pH 10 at 4°C). Fractions were analyzed by SDS-PAGE, quantified on a NanoDrop spectrometer (Thermo Scientific), and stored at -80°C.

### *In vitro* transcription

RNA transcripts for *in vitro* splicing, EMSAs, and UV-crosslinking were transcribed from pGEM vectors with T7 RNA polymerase as described previously (5, 14, 15), albeit to aid in visualization, [α-^32^P]CTP was used to label the RNAs instead of [α-^32^P]UTP.

For high yield RNA transcripts for splicing assays and EMSAs, the reaction contained 1 μg of linearized DNA, 2 μL of 5x transcription buffer (30mM MgCl_2_, 10 mM spermidine, 0.2 M Tris-HCl, pH 7.5), 1μL 10x rNTPs (5 mM UTP, 5 mM ATP, 5 mM GTP, 2 mM CTP), 0.5 μL RNasin (Promega), 0.5 μL 10 mM RNA cap analogue (Promega), 2.5 μL ddH_2_O, 1.5 μL T7 RNA polymerase, and 1 μL of 0.37 MBq/μL [α-^32^P]CTP. The transcripts for UV-crosslinking required high specific activity and the reactions were adjusted to 0.4 mM CTP and 2 μL of 0.37 MBq/μL [α-^32^P]CTP. Reactions were incubated for 2 hours at 37°C followed by a 1:5 dilution with ddH_2_O and 1 μL was taken for total scintillation counts. RNAs were then phenol extracted and spun in a G-50 column (ProbeQuant) at 3,000 x g for 2 min to purify the RNAs. A Beckman LS 3801 scintillation counter was used to quantify the radioactivity of 1 μL of purified RNAs along with the 1 μL total sample. The percentage yield was calculated by dividing the [incorporated RNA counts x final volume] by [total RNA counts x 50 μL]. The [percent yield/100] x [mass of the nucleotide x number of nucleotides x CTP concentration x volume] was used to calculate the amount of RNA transcribed. The transcribed RNAs were then diluted to 20 fmol/μL with ddH_2_O. The transcripts used in the splicing assays contained α-tropomyosin exons 1, 3, and 4 and the introns between exons 1-3 and 3-4 (known as TM1-3-4) providing a sufficient pre-mRNA transcript to analyze splicing activity. The transcripts used in the UV-crosslinking assays were a truncated version of TM1-3-4, containing only the exon 3 and flanking introns. While the transcript used in the EMSAs was a synthetic RNA of 120 nucleotides containing three tandem CAC motifs modeled after the intronic region between exons 1 and 3 of TM1-3-4 (Yi Yang et al., manuscript in preparation).

### *In vitro* splicing assay

Splicing reactions contained 10 μL total volume: 2.6% polyvinyl alcohol, 2 mM MgCl_2_, 500 μM ATP, 20 mM creatine phosphate, 300 U/μL RNasin (Promega), 12 mM Hepes, pH 7.9, 0.3 mM DTT, 30% HeLa nuclear extract, and 12.5 ng α^32^P labelled pre-mRNA. Reactions were incubated at 30°C for 3 hours, subjected to proteinase K digestion, phenol extracted, and ethanol precipitated. Reaction products were finally resolved on an 8 M urea 4% polyacrylamide gel. Gels were then dried and visualized by autoradiography on a GE Typhoon FLA 9000. Nuclear extracts were prepared from HeLa cells in suspension using the modifications described previously (16).

### Electrophoretic mobility shift assay

Wild-type recombinant RBPMS-A or the mutant variants were titrated (4, 2, 0.66, 0.22, 0.07, 0.024, 0 μM) and incubated with 20 fmol of *in vitro* transcribed RNA in binding buffer (50 mM HEPES pH 7.2-7.9, 15 mM MgCl_2_, 25% glycerol, 5 mM DTT, 45 mM KCl, 5μg tRNA) for 25 minutes at 30°C. Heparin (5 μg) was then added to remove non-specific binding and incubated for an additional 5 minutes at 30°C. 1 μL of 50% glycerol was added to the reactions before loading onto a 4% 60:1 acrylamide:bisacrylamide non-denaturing gel with TBE running buffer at room temperature. Gels were then dried and visualized by autoradiography on a GE Typhoon FLA 9000.

### UV-crosslinking

The same binding reaction described above for EMSAs was used for UV-crosslinking, however, 20 fmol of the truncated transcript was used with or without 20% HeLa nuclear extract. Reactions were then UV-crosslinked on ice in a Stratalinker with two pulses at 960 mJ. An RNase mix (RNase A, 12.5 μg and RNase T1, 0.35 U) was added and the reactions were incubated for 10 minutes at 37°C. Reactions were quenched by the addition of 4x SDS sample buffer, boiled at 80°C for 5 minutes, and loaded onto a 15% SDS-PAGE gel. Gels were then dried and visualized by autoradiography on a GE Typhoon FLA 9000.

### *In vitro* kinase assay

Kinase assays contained wild-type recombinant RBPMS-A or the non-phosphomimetic T/A mutant and active recombinant ERK2 (obtained from Remkes Scheele, Department of Biochemistry, University of Cambridge). Myelin Basic Protein (MBP; Millipore) was used as a positive control. Kinase reactions were 30 μL total: 3 μL 10x kinase buffer (400 mM Tris-HCl pH 7.5,100 mM MgCl_2_, 1 mM EGTA, 10 mM DTT, 1x phosphatase inhibitor), 1 μL kinase (various concentrations), 1 μL substrate (various concentrations), 23 μL ddH_2_O, and 2 μL ATP mix (0.5 mM ATP, 0.74 MBq [γ-^32^P]ATP – for radiolabeled assays). Reactions were incubated for 5 minutes at room temperature and quenched by the addition of 4x SDS sample buffer. Samples were separated by 15% SDS-PAGE. Gels were then dried and visualized by autoradiography on a GE Typhoon FLA 9000.

### Analytical Ultracentrifugation

An Optima XL-I (Beckman Coulter) centrifuge using an An60 Ti eight-hole rotor was used to measure sedimentation velocity (SV). Standard 12 mm double-sector Epon centerpieces with sapphire windows contained 400 μL of wild-type recombinant RBPMS-A or the mutant variants that were buffer exchanged with ultracentrifugation buffer (20 mM HEPES, 500 mM KCl, 1 mM TCEP, pH adjusted to 7.9) and adjusted to 0.5 mg/mL. Interference data were acquired at 40,000 rpm, 20°C, overnight. 10 scans were overlaid as a single point to reduce file size. The partial specific volume of the protein (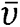 = 0.73 mL/g) and the viscosity and density of the buffer (η = 1.01036 x 10^-2^ mPa·s; ρ = 1.0232 g/mL) were calculated using SEDNTERP (Laue et al., 1992). 100 scans taken over 10 hours were used to determine the multi-component sedimentation coefficient c(s) distributions by direct boundary modeling of the Lamm equation using SEDFIT (v14.1) (17). The proportion of dimers to oligomers was determined by integration of the peaks, which were identified by their fitted mass.

### Phosphatase Assay

Whole-cell extracts of HEK293T cells overexpressing our protein of interest (see transient transfection in methods) were harvested in gentle lysis buffer (150mM NaCl, 1% Triton X-100, 0.1% SDS, 50mM Tris pH 8). Extracts (3μL) were incubated in NEBuffer Pack for Protein MetalloPhosphatases (PMP; 50mM HEPES, 100mM NaCl, 2 mM DTT, 0.01% Brij 35, pH 7.5), 10 mM MnCl_2_, and 4 U of lambda protein phosphatase for 30 min at 30°C. Protein was then resolved on a 15-20% SDS-PAGE followed by immunoblotting.

### Immunoprecipitation

Wild-type pCI-3xFLAG-RBPMS-A or pCI-3xFLAG-RBPMS-A ΔF-Site along with pCI-3xFLAG-MEK1-CA and pCI-3xFLAG-ERK2 were transiently transfected into HEK293T cells for 36 hours. Whole-cell extracts were harvested in 1 mL (per 10 cm dish; see seeding density in transient transfection method) of M-PER^TM^ Mammalian Protein Extraction Reagent plus 40uL 25x cOmplete^TM^ Protease Inhibitor Cocktail (Sigma-Aldrich 4693132001) and processed according to manufacturer’s guidelines. 100 μL of cell lysates was added to 900 μL of ice-cold NETS Buffer (150 mM NaCl, 50 mM Tris pH 7.5, 5mM EDTA, 0.05% NP40) along with rabbit anti-RBPMS antibody (1:500) or IgG control antibody (1:500, SMPTB pre-immune 1958). After a rotating incubation for 60 mins at 4°C, Protein A magnetic beads were added and incubation continued for an additional 60 mins at 4°C, rotating. Antibodies bound to the protein of interest along with any other protein-protein interactions were pulled down with a magnetic separation rack. Proteins were resolved on a 20% SDS-PAGE followed by immunoblotting.

### NMR spectroscopy

For expression of ^15^N- or ^13^C, ^15^N-labelled RBPMS and mutant proteins, ^13^C_6_-glucose and/or ^15^NH_4_Cl were used as the sole carbon/nitrogen source in M9 minimal medium. NMR measurements were made on ∼ 0.5 mM solutions in 10% ^2^H_2_O, 10 mM HEPES (pH 6.8) and 50 mM KCl. Experiments were recorded on Bruker AVANCE III 600 or 800 MHz spectrometers equipped with QCI or TXI cryoprobes. Experiments were recorded at both 37°C (optimal for the folded regions) and 5°C (optimal for the disordered tails). Established versions (18) of HNCA, HN(CO)CA, HN(CO)CACB, HNCACB, HNCO, 3D TOCSY-^15^N-HSQC, and NOESY-^15^N-HSQC experiments were acquired, alongside HNN (19). Triple resonance experiments were acquired with 25% nonuniform sampling, using Poisson-gap sampling (20), and reconstructed using the Cambridge CS package and the CS-IHT algorithm (21). Data were processed using the AZARA suite of programs (v. 2.8, © 1993–2022; Wayne Boucher and Department of Biochemistry, University of Cambridge, unpublished). Backbone assignments were available for a related protein (BMRB Entry 19298, (22)), which were used as a starting point. Assignment was carried out using CcpNmr Analysis v. 2.4 (23). Chemical-shift differences were calculated using Δδ = [(Δδ^H^)^2^ + (0.15 × Δδ^N^)^2^]^1/2^ (24).

### SEC-MALS

The absolute molecular masses of the truncated RBPMS-A WT and T/E protein samples (used for NMR) were determined by SEC-MALS. 50-μl protein samples (at approximately 2 mg/ml) were loaded onto a Superdex 75 10/300 GL increase size-exclusion chromatography column (GE Healthcare) in 10 mM HEPES, pH 6.8, 50 mM KCl, at 0.5 ml/min with an ÄKTA Purifier (GE Healthcare). The column output was fed into a DAWN HELEOS II MALS detector (Wyatt Technology) followed by an Optilab T-rEX differential refractometer (Wyatt Technology). Light scattering (LS) and differential refractive index (dRI) data were collected and analyzed with ASTRA 6 software (Wyatt Technology). Molecular masses and estimated errors were calculated across individual eluted peaks by extrapolation from Zimm plots with a dn/dc value of 0.1850 ml g−1. SEC-MALS data are presented with LS and UV plotted alongside fitted molecular masses.

## RESULTS

### Phosphorylation of RBPMS at Thr113 and Thr118

Post-translation modifications (PTMs) of RBPMS were determined using the PhosphoSitePlus database which collates PTMs of proteins from published datasets (25), filtering for entries from high throughput papers supported by five or more references. Two strongly supported events, phosphorylation of Thr113 and Thr118, occur immediately on the C-terminal side of the RBPMS RRM domain (Figure 1A; Supplementary Figure S1A, original data from refs (26–30)). Not only were these residues and their adjacent prolines phylogenetically conserved (Thr113 across vertebrates, and Thr118 across vertebrates except fish), but phosphorylation of Thr113 and Thr118 was found in human, rat, and mouse datasets (Supplementary Figure S1B). Moreover, the two phosphorylation events frequently occur together, and in one study were identified in M phase arrested HeLa cells (26)(27). Taken together, these data suggest that the possible canonical phosphorylation of RBPMS is at residues T113 and T118, and that phosphorylation of RBPMS could perhaps occur in a cell-cycle-specific fashion.

### Phosphomimetic mutations inhibit RBPMS splicing activity in cell culture

Since the most likely phosphorylation of RBPMS occurs on residues Thr113 and Thr118, we made mutations to these residues in the canonical RBPMS-A isoform. We created both non-phosphomimetic and phosphomimetic mutants by mutating both threonine 113 and 118 to either alanine in “RBPMS-A T/A” or glutamic acid in “RBPMS-A T/E”, as well as the individual mutants, T113E and T118E. Mutants were transiently transfected into HEK293T cells, and their splicing activity was evaluated by assessing their ability to promote SMC AS of endogenous *ACTN1, TPM1, FLNB*, and *MPRIP* (Figure 1B and D). *ACTN1* and *TPM1* contain non-smooth muscle exons (NM), exons found in the more motile and proliferative phenotype, that are repressed by RBPMS-A thereby allowing inclusion of the mutually exclusive smooth muscle (SM) exon, while *FLNB* and *MPRIP* contain SM cassette exons that are activated by RBPMS-A (5). While WT RBPMS promoted the SMC splicing pattern in each case (Fig 1B, lanes 1 and 2), the phosphomimetic mutant, RBPMS-A T/E, had significantly reduced activity across all transcripts (Figure 1B and D). RBPMS-A T/E significantly failed to repress the NM exons of *ACTN1* and *TPM1* and had significantly impaired activation of the SM exons of *FLNB* and *MPRIP* when compared to wild-type (WT) (Figure 1B, lane 4 vs 2). However, the non-phosphomimetic, RBPMS-A T/A, retained similar repressive and activation activity compared to WT RBPMS-A (Figure 1B, lane 3 vs 2). All proteins were expressed at similar levels (Figure 1C), thus the reduced activity of the phosphomimetic can be directly attributed to the phosphomimetic mutations. Although the single T113E and T118E mutations showed some reduced splicing activity for some transcripts, neither of them had a significant reduction in activity across all transcripts, which was only observed for the double mutant (Figure 1B and D).

We next set out to determine the effects of the phosphomimetic mutations on RBPMS activity in PAC1 vascular smooth muscle cells. Due to the poor transfection efficiency of plasmids in PAC1 cells, we generated doxycycline-inducible cells using the pInducer22 lentiviral vector (13) to express our RBPMS-A wild-type and mutant constructs. Following a 24-hour doxycycline induction, RNA was harvested, and the splicing activity was evaluated by assessing the splice variants of endogenous *Actn1, Tpm1, Flnb*, and *Ptprf* (Figure 1E and G). As observed in the HEK293T cells, the WT and T/A mutant RBPMS regulated all splicing events (lanes 4 vs 3 and 6 vs 5), but the RBPMS-A T/E mutant failed to repress the NM exons in *Actn1* and *Tpm1* and was significantly impaired for activation of the SM exons in *Flnb* and *Ptprf* (Figure 1E, lane 8 vs 4). Similar expression levels of all RBPMS-A proteins were observed (Figure 1F). Collectively, these data from the HEK29T transfections and now the PAC1 overexpression show that phosphomimetic mutations of RBPMS-A at T113 and T118, and by extension potentially phosphorylation, down-regulates both its exon repression and exon activation splicing activity.

To ensure that the phosphomimetics were not disrupting any regulatory elements that might exist around the area of the mutations, we deleted residues 112 through 119 (Δ112-119) to capture the region where T113 and T118 reside (Supplementary Figure S1C). This mutant was then transiently transfected into HEK293T cells, and its splicing activity was evaluated by assessing the splice variants of endogenous *ACTN1, TPM1, FLNB*, and *MPRIP* (Supplementary Figure 1E and F). There was no significant change in splicing activity across all transcripts as the deletion mutant had similar activity to the wild-type protein (Figure 1D). These data suggest that the region containing T113 and T118 is not essential for RBPMS-A activity but acts purely as a site for regulatory input by phosphorylation.

### RBPMS-A is a substrate of ERK2

Since the phosphomimetic T/E mutant suggested that the splicing activity of RBPMS-A was down-regulated by its phosphorylation, we wanted to identify the kinase(s) responsible for phosphorylation at T113 and T118. First, we used PhosphoNET Kinase Predicter (www.phosphonet.ca), which determined that the most likely kinase candidate for RBPMS at T113 and T118 was ERK2, with ERK1 identified as the second most likely candidate (Figure 2A). Secondly, we searched current literature for known ERK substrates and found that mouse RBPMS has been shown to be phosphorylated by ERK2 at T118 (31). Mouse RBPMS shares a 98% identity with human RBPMS and is conserved at T113 and T118 (Supplementary Figure S1B). Finally, growth factors, such as PDGF, drive the dedifferentiation of VSMCs by activating the Ras/Raf/MEK/ERK pathway, which could rapidly inactivate the RBPMS-driven splicing program (3).

**Figure 2.**
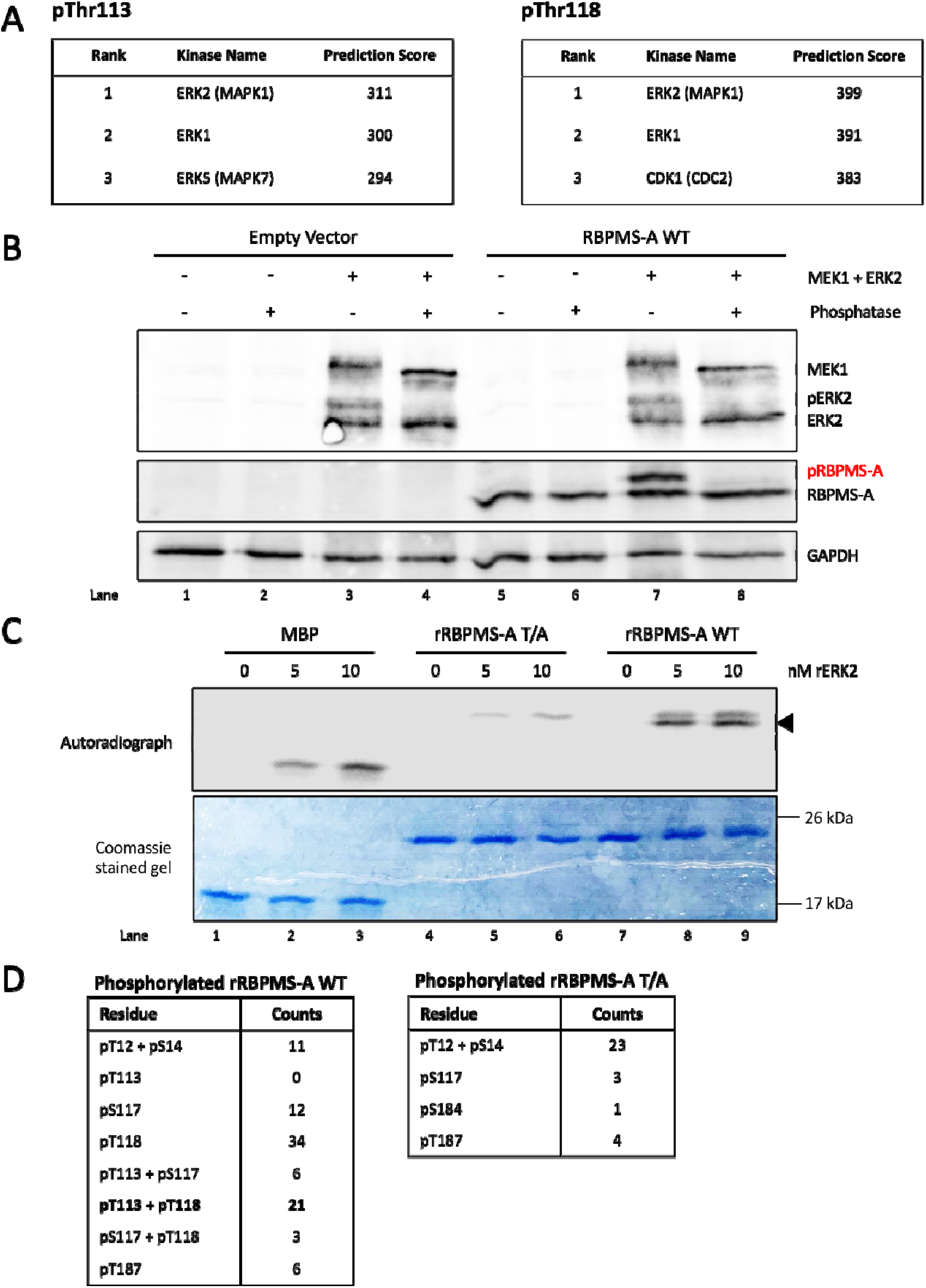
ERK2 Phosphorylates RBPMS-A. (A) Top 3 predicted kinase scores for RBPMS-A phosphorylation at either Thr113 or Thr118 were obtained using Kinase Predictor V2 on PhoshoNET (www.phosphonet.ca). Scores were calculated accounting for the presence of neighboring phosphosites. Scores are not scaled to a maximum value. (B) Immunoblot of transiently transfected wild-type (WT) RBPMS-A with (+) or without (-) constitutively active (CA) MEK1 and wild-type ERK2. Following protein harvest, whole-cell lysates were treated (+) or mock-treated (-) with lambda phosphatase. Phosphorylated RBPMS-A (pRBPMS-A) is highlighted in red. Activated (phosphorylated) ERK2 (pERK2) can be seen in lanes 3 & 7. Immunoblot was probed with antibodies to FLAG and GAPDH. GAPDH was used as a loading control. (C) Recombinant wild-type (WT) RBPMS-A or RBPMS-A T/A were incubated with increasing nM amounts of highly active recombinant ERK2 in the presence of [γ-^32^P]ATP in an *in vitro* kinase assay. Proteins were stained with Coomassie blue (bottom) followed by autoradiography (top). Phosphorylation of RBPMS-A WT at T113 and T118 is indicated by the black arrow. Myelin Basic Protein (MBP) was used as a positive control. (D) Mass spectrometry results of *in vitro* kinase assay for either phosphorylated RBPMS-A WT or the T/A mutant. Phosphorylation sites of interest are bolded.

Given this information, we set out to determine if RBPMS could be phosphorylated by ERK2. A phosphorylation-dependent mobility shift occurs for many eukaryotic proteins in SDS-PAGE (32); therefore, to observe the phosphorylation of RBPMS-A in a cell culture model, we transiently transfected wild-type RBPMS-A along with constitutively active MEK1 (MEK1-CA) and wild-type ERK2 in HEK293T cells and analyzed the protein via western blot. The results showed substantial phosphorylation of RBPMS-A in the presence of MEK1-CA and ERK2 as a large proportion of RBPMS-A had shifted upwards indicating phosphorylation (Figure 2B, lanes 7 vs 5). To confirm bona fide phosphorylation of RBPMS-A, we carried out lambda phosphatase treatment prior to western blotting. The collapse back to a single lower RBPMS-A band confirms that the upper band corresponds to phosphorylated RBPMS-A (Figure 2B, lane 8).

Next recombinant wild-type RBPMS-A was incubated *in vitro* with recombinant ERK2 and [γ-^32^P] ATP. Myelin basic protein (MBP), a known target of ERK2, was used as a positive control. RBPMS-A WT showed two radiolabeled bands (Figure 2C, upper panel, lanes 8 and 9) indicating that ERK2 can phosphorylate wild-type RBPMS-A at two or more positions. The RBPMS-A T/A showed only the fainter lower mobility band (lanes 5 and 6). The higher intensity lower bands (Figure 2C, arrow) are present only in the wild-type RBPMS-A lanes suggesting that these correspond to phosphorylation at T113 and/or T118. The upper bands present in the wild-type RBPMS-A lanes and RBPMS-A T/A lanes presumably correspond to phosphorylation of the protein at additional sites. Indeed, in addition to T113/118, both T12 and S144 also lie within X[pS/pT]P and PXxx[pS/pT]P consensus contexts for phosphorylation by ERK2 (33, 34). Together, these results demonstrate that RBPMS-A is a substrate of ERK2.

### Identification of RBPMS-A sites phosphorylated by ERK2

To directly determine which residues are phosphorylated by ERK2, we performed an in vitro kinase assay with unlabeled ATP, followed by mass spectrometry. The bands sent for analysis are shown in Supplementary Figure S2A. The results indicated that ERK2 phosphorylates RBPMS-A at Thr12, Ser14, Thr113, Ser117, Thr118, and Thr187 (Figure 2D), with the overwhelming majority of peptides corresponding either to single phosphorylation at Thr118 or dual phosphorylation at Thr113 and Thr118. Of note, phosphorylated Thr113 was only observed in conjunction with phospho-Thr118, suggesting that phosphorylation of Thr118 is necessary before phosphorylation occurs at Thr113, as previously reported (26, 27).

Since Ser14 and Thr187 do not lie within a consensus context for ERK2 phosphorylation and their phosphorylation was observed at a lower frequency than Thr113 and 118, we concluded that these phosphorylation events might be due to the lower specificity of *in vitro* kinase assays so we did not consider them further (35). Although Ser144 has the sequence context for phosphorylation by ERK2, peptides containing S144 were not detected by mass spectrometry after various protease digestions, although subsequent experiments (below) indicated that S144 is phosphorylated by ERK2 in cells. Transfected phosphomimetic and non-phosphorylatable mutants at Thr12 and Ser144 (T12E, T12A, S144E, S144A, Figure 3A, B) and Ser 117 (S117E, Supplementary Fig S3) had identical splicing activity to WT RBPMS, indicating that phosphorylation at these residues plays no role in regulating RBPMS splicing activity. However, Thr12 and Ser144 and their adjacent prolines are conserved across vertebrates, suggesting that their phosphorylation may play some other role.

**Figure 3.**
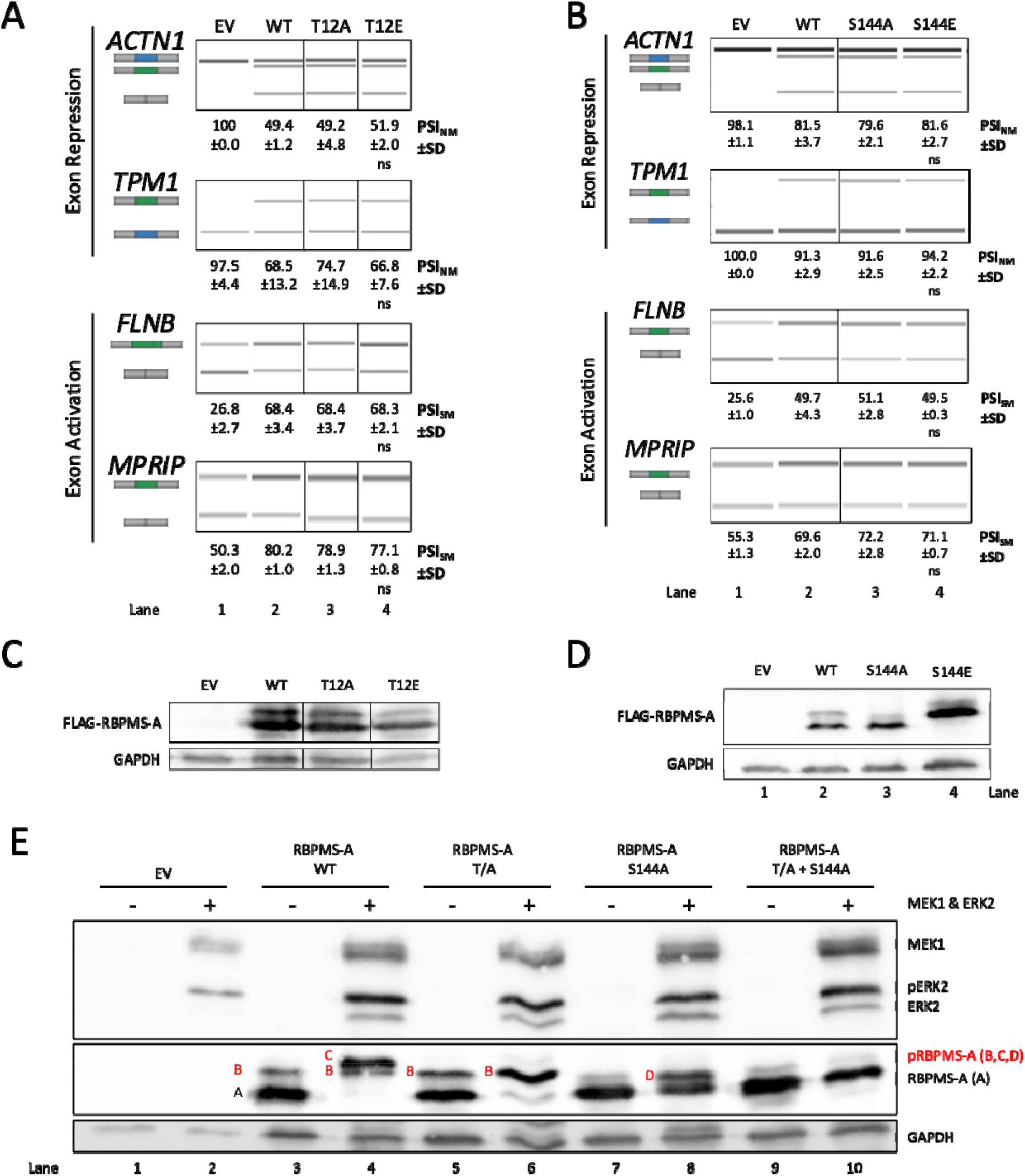
Other Sites of Phosphorylation Identified on RBPMS-A Have No Effect on Splicing. (A&B) Splicing analysis of the exon repression function of wild-type (WT) RBPMS-A and effectors in endogenous *ACTN1* and *TPM1*, and its exon activation function in endogenous *FLNB* and *MPRIP* in HEK293T cells. RT-PCR followed by visualization on QIAxcel. Cartoons below the gene names represent the PCR products. The blue exon in the cartoon represents the NM (Non-smooth Muscle) exon, while the green exon represents the SM (Smooth Muscle) exon. PSI (Percent Spliced In) values are the mean ± SD (n = 3). A Student’s t-test was used to determine statistical significance between wild-type RBPMS-A and effector. EV=empty vector. (C&D) Immunoblot of FLAG-RBPMS-A effectors used for splicing assays in (A&B). Whole-cell lysates were probed with antibodies to FLAG and GAPDH. GAPDH was used as a loading control. (E) Immunoblot of transiently transfected wild-type (WT) RBPMS-A and mutants with (+) or without (-) constitutively active (CA) MEK1 and wild-type ERK2. Phosphorylated RBPMS-A (pRBPMS-A) is highlighted in red. A=unphosphorylated RBPMS-A, B=pS144, C=pS144+pT113+pT118, D=pT113+pT118. Activated (phosphorylated) ERK2 (pERK2) can be seen in lanes 2, 4, 6, 8 & 10. Immunoblot was probed with antibodies to FLAG and GAPDH. GAPDH was used as a loading control.

Cotransfection experiments using non-phosphorylatable mutants T113/118A (T/A), S144A, and a combined triple mutant T/A+S144A, confirmed that these three residues are phosphorylated by ERK2 in HEK293T cells (Figure 3E). Two WT RBPMS bands were observed in the absence of MEK-CA and ERK2 (Fig 3E, lane 3) corresponding to non-phosphorylated RBPMS (band “A”) and RBPMS basally phosphorylated at S144 (band “B”). Upon addition of MEK-CA and ERK2 (lane 4), band A disappeared and a band corresponding to phosphorylation at T113/T118 and S144 appeared (band “C”), with some residual band B. The identity of bands B and C was confirmed by the the non-phosphorylatable mutants. The T/A mutant appeared identical to WT under resting conditions (lanes 3 and 5) but upon the addition of MEK-CA and ERK2, band B became more intense, and no band C was observed. With S144A, band B was not present, consistent with it corresponding to phosphorylation at S144 (lane 7). An additional band with mobility intermediate between A and B increased upon the addition of MEK1-CA and ERK2 (Figure 3E, lanes 7 and 8, band D), corresponding to phosphorylation at T113 and T118. For the triple mutant (T/A + S144A) bands B, C, and D were completely abolished (Figure 3E, lane 10) consistent with their identification as RBPMS phosphorylated at S144, T113/T118/S144 and T113/T118 respectively.

Many kinases are recruited by specific docking sites located on their substrates. The motif 167-<u>F</u>T<u>YP</u>-170 on RBPMS is a perfect match to the ERK2 F-site docking motif FX(F/Y)P, and is located in a canonical position downstream of the phosphorylation sites (33, 36–40). Consistent with its identification as an F-site, ERK2 was observed to coimmunoprecipitate with RBPMS-A (Figure 4B), and this was reduced approximately 5-fold upon deletion of RBPMS aa 167-170 (Figure 4B, ΔF-site). This result indicated that 167-<u>F</u>T<u>YP</u>-170 is important for ERK2 binding. Phosphorylation by ERK2 was also substantially reduced for the F-site mutant compared to WT RBPMS (Figure 4C, lane 4 vs 2). The basal level of S144 phosphorylation of WT and T/A RBPMS-A was undisturbed by the F-site deletion (Figure 4C, lanes 3 and 7 compared to 1). However, deletion of the F-site reduced the level of ERK2 induced S144 phosphorylation in the T/A mutant (Figure 4C, lanes 6 and 8), while phosphorylation of Thr113 & Thr118 appears to be reduced by the removal of the F-site in the S144A mutant (Figure 4C, lanes 10 and 12). Together, these data suggest that the RBPMS-A F-site is important for ERK2 binding and phosphorylation of Thr113, 118 and Ser144. In contrast, F-site deletion had no effect on splicing activity (Supplementary Figure S4A, B, C), similar to the effect of deleting the region where T113 and T118 reside. Taken together, the preceding data indicate that ERK2 phosphorylates RBPMS at T113, T118, and S144, but that regulation of splicing activity results only from phosphorylation of RBPMS-A at both Thr113 and Thr118.

**Figure 4.**
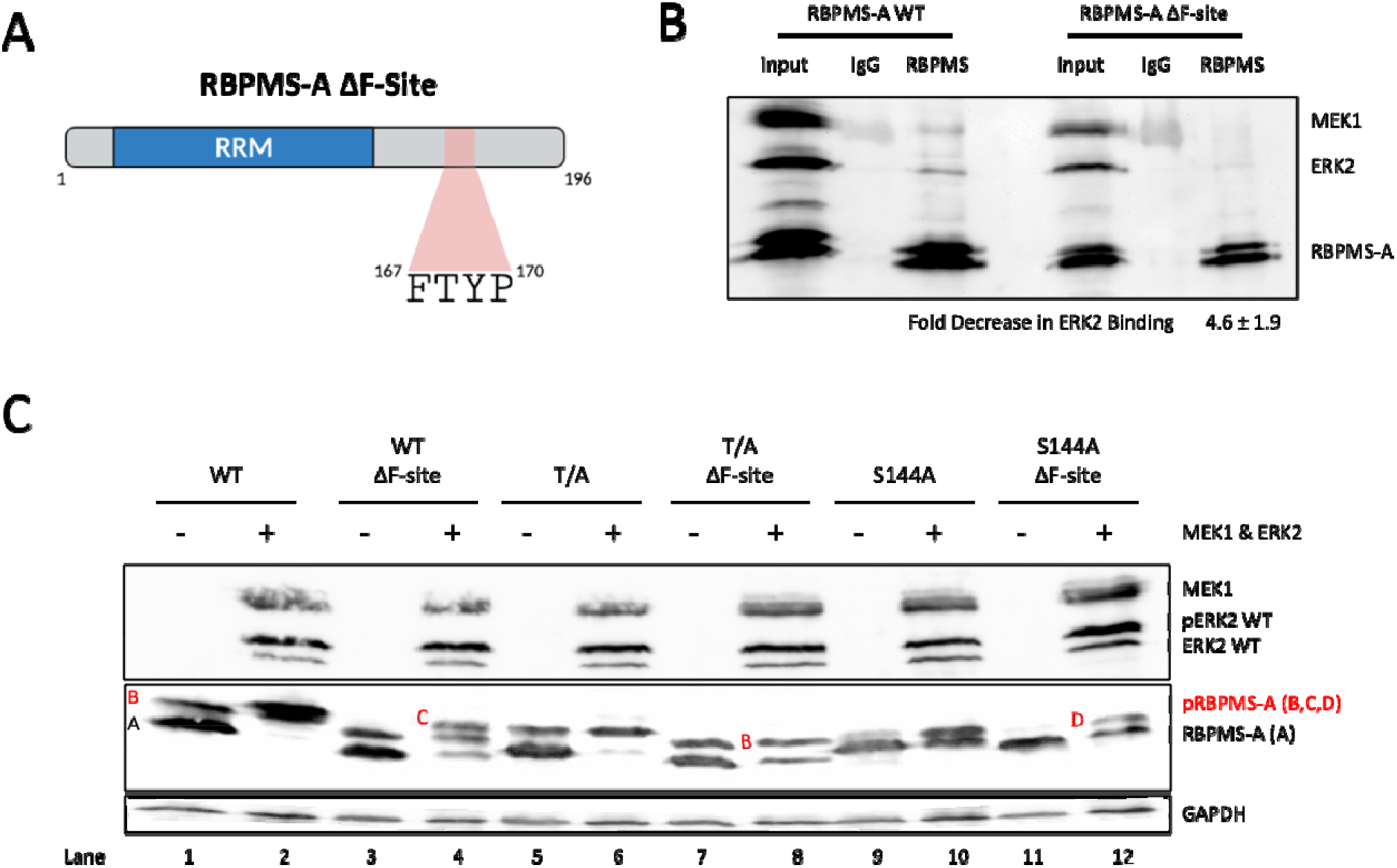
ERK2 Binds to F-site Located in RBPMS-A C-terminal Tail. (A) Graphical representation of RBPMS-A ΔF-Site with the F-site region highlighted in red. The ΔF-Site mutation is the deletion of FTYP. Created with BioRender.com (B) Confirmation of ERK2 and RBPMS-A binding. HEK293T cell lysates expressing either FLAG-RBPMS-A WT or the F-site mutant (FLAG-RBPMS-A ΔF-Site) along with FLAG-MEK1 and FLAG-ERK2 was precipitated with anti-FLAG. Fold decrease in ERK2 binding to RBPMS was normalized to immunoprecipitated RBPMS-A (n=3), the value is the mean ± SEM. (C) Immunoblot of transiently transfected wild-type (WT) RBPMS-A and mutants with (+) or without (-) constitutively active (CA) MEK1 and wild-type ERK2. Phosphorylated RBPMS-A (pRBPMS-A) is highlighted in red. A=unphosphorylated RBPMS-A, B=pS144, C=pS144+pT113+pT118, D=pT113+pT118. Activated (phosphorylated) ERK2 (pERK2) can be seen in lanes 2, 4, 6, 8, 10, & 12. Immunoblot was probed with antibodies to FLAG and GAPDH. GAPDH was used as a loading control.

### Phosphomimetic mutations directly impair RBPMS-A splicing activity

One way in which phosphorylation can influence splicing factor activity is by altering localization (41). To determine if the subcellular localization of RBPMS-A was affected by mutations in T113/118. FLAG-tagged RBPMS-A wild-type and the T/A and T/E mutants were transiently transfected in HEK293T cells and subcellular localization was analyzed by immunofluorescence microscopy. Both wild-type and T/A mutant RBPMS-A were localized primarily in the nucleus but excluded from nucleoli. In contrast, the T/E mutant was present in both the nucleus and the cytoplasm (Figure 5A). In an attempt to address whether the reduced splicing activity of the RBPMS-A T/E mutant was due to its partial re-localization to the cytoplasm, we added an N-terminal exogenous nuclear localization signal (NLS) to each of the constructs (WT, T/A, T/E) and transiently transfected them into HEK293T cells to evaluate their splicing activity (Supplementary Figure S5A and B). Surprisingly, the exogenous NLS failed to rescue the partial cytoplasmic localization (Supplementary Figure S5D) of the RBPMS-A T/E mutant and also had no effect on splicing activity (Supplementary Figure S5A and B).

**Figure 5.**
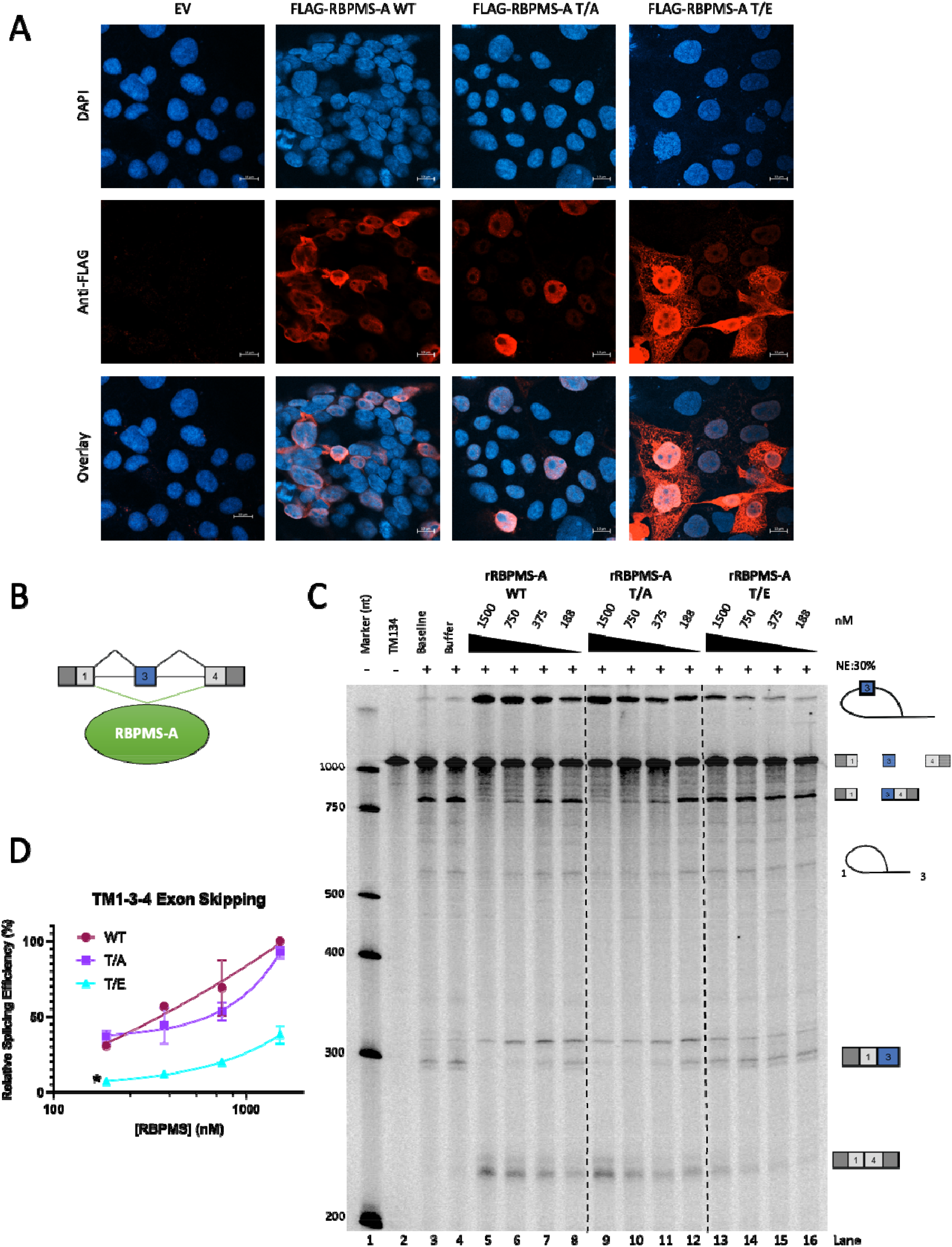
Loss of RBPMS-A Splicing Activity is Independent of Localization. (A) Immunofluorescence microscopy images of HEK293T cells overexpressing wild-type (WT) RBPMS-A and the T/A and T/E mutants. All images were taken with the same parameters using a 40X/1.20W Korr objective. Top row shows DAPI staining for DNA, middle row shows staining for FLAG tagged effectors, and bottom row is an overlay of the DAPI and FLAG images. Scale bars are 10uM. EV = empty vector. (B) Graphical representation of the full-length TM1-3-4 pre-mRNA showing that RBPMS-A represses exon 3 (blue). (C) *in vitro* splicing of TM1-3-4 RNA by recombinant wild-type (WT) RBPMS-A and the T/A and T/E mutants in 30% HeLa nuclear extracts (NE) indicated by the (+). A decreasing 2-fold dilution series of recombinant protein was used as indicated by the black triangles and the nM amounts listed above. “TM134” is the RNA alone. “Baseline” is TM134 with the addition of the NE buffer. “Buffer” is TM134 with the addition of the splicing buffer. The cartoons on the right represent all the splicing intermediates and products. (D) Quantification of the relative TM1-3-4 exon skipping or 1-4 product found in (C) of each effector normalized to wild-type RBPMS-A after background splicing in the buffer was taken into consideration. An ordinary one-way ANOVA was used to determine statistical significance between wild-type RBPMS-A and effector across all concentrations, indicated by *p < 0.05. Error bars specify SD (n = 3).

The immunofluorescence data indicated that only a fraction of the phosphomimetic T/E mutant is re-localized to the cytoplasm, which might not be sufficient to account for the loss of splicing regulation. To directly assess whether the phosphorylation of RBPMS-A directly affects its splicing activity, independent of localization, we expressed and purified recombinant RBPMS-A wild type, T/A, and T/E (Supplementary Figure S6A). Previous work in the lab has shown that wild-type rRBPMS-A represses (skips) exon 3 of a TM1-3-4 transcript in cell-free splicing assays, dependent on flanking CAC motifs (Yi Yang et al., manuscript in preparation; Figure 5B). Therefore, using the well-characterized TM1-3-4 minigene (15, 42), *in vitro* splicing assays were carried out with the recombinant RBPMS proteins to evaluate their splicing activity (Figure 5C). In the absence of added RBPMS, the 1-3-4 transcript primarily splices to include exon 3, indicated by the fully spliced 134 product as well as the prominent 1-34 partially spliced RNA (lanes 3 and 4). Upon addition of rRBPMS-A WT, a switch in splicing was observed with dose-dependent decreases in exon 3 inclusion bands and a reciprocal increase in bands associated with exon skipping, the spliced 14 product, and the corresponding lariat that contains exon 3 as part of the intron (lanes 5-8 compared to lane 4). The rRBPMS-A T/A mutant showed similar activity to WT (lanes 9-12 compared to lane 4), as in cell transfection assays. However, in agreement with our observations in HEK293T and PAC1 cells, the rRBPMS-A T/E mutant showed near-complete loss of activity (lanes 13-16 compared to lane 4); at the highest concentration of T/E protein a slight increase in exon skipping products was observed, but exon 3 inclusion products were unaffected. These data demonstrate that the T113/118 phosphomimetic mutation directly reduces its splicing activity, suggesting that the loss of activity in cells (Figure 1B and E) is not merely localization-dependent.

### Phosphomimetic mutation of RBPMS-A results in reduced RNA-binding affinity

To determine how the phosphorylation of RBPMS-A affects its splicing activity, we tested the effects of T113/118 mutations (T/E) upon RNA-binding. RBPMS dimers bind to tandem CAC motifs separated by a variable nucleotide spacer (6, 43), and we have shown that RBPMS-A can bind to the three dual CAC motifs found in the intron upstream of *Tpm1* exon 3 (5). Therefore, we *in vitro* transcribed the intronic region upstream of *Tpm1* exon 3 (see Methods) for use in an electrophoretic mobility shift assay (EMSA) to examine the binding of rRBPMS-A WT and mutant proteins (Figure 6A). By estimating the concentration of protein at which 50% of RNA is bound we determined the apparent dissociation constants for comparison between the wild-type and mutants. The rRBPMS-A T/E mutant has a higher apparent dissociation constant (K_d_ = 756.1 nM) compared to wild-type (K_d_ = 212.9 nM) and the T/A mutant (K_d_ = 267.9 nM) (Figure 6B). To further explore how phosphorylation affects RNA-binding, UV-crosslinking assays were carried out in the presence and absence of HeLa nuclear extract using the same *Tpm1* RNA (Figure 6C). Under both conditions, the RBPMS-A T/E mutant showed a significant reduction in crosslinking efficiency compared to WT. The reduction was more substantial in the presence of nuclear extract with only 7.8% crosslinked compared to WT (at 4 uM; Figure 6C, upper panel). Thus, although the T/E mutant only had a 3-fold lower affinity for the *Tpm1* RNA (Figure 6A,B), in the presence of other RNA-binding proteins in nuclear extract the lower affinity leads to substantially lower RNA-binding. Taken together, these data suggest that phosphomimetic mutant (and by extension phosphorylation) of RBPMS-A inhibits RNA-binding, thereby explaining the loss of splicing regulatory activity.

**Figure 6.**
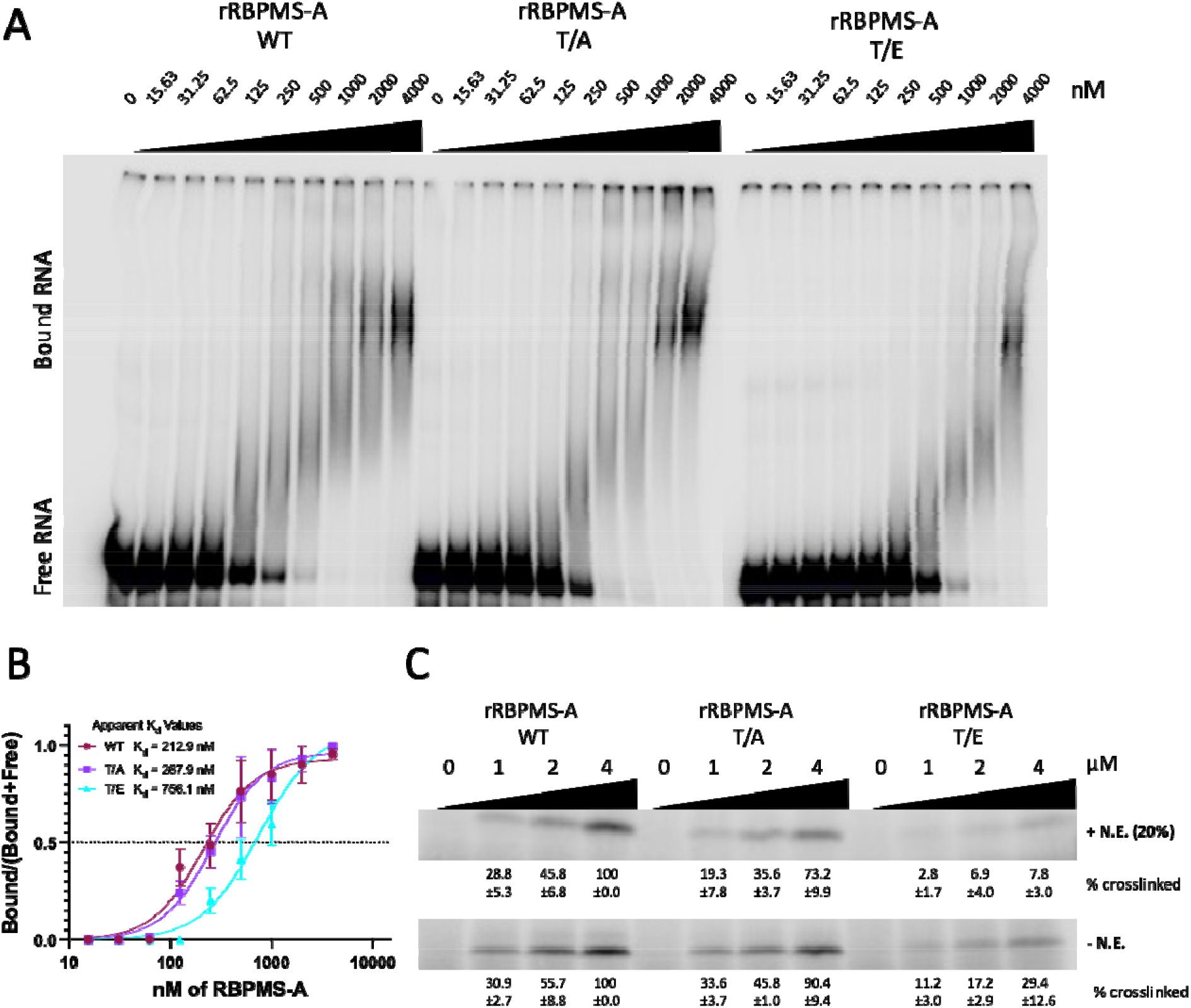
RBPMS-A Phosphomimetic Fails to Bind to RNA. (A) Electrophoretic Mobility Shift Assays (EMSAs) using recombinant wild-type (WT) RBPMS-A and the T/A and T/E mutants and an *in vitro* transcribed intron upstream of *Tpm1* exon 3. A decreasing serial dilution (1:10) of recombinant protein was used as indicated by the black triangles and the nM amounts listed above. Unbound or free RNA can be found at the bottom of the gel and the RBPMS + RNA complexes can be seen towards the upper right side of each gel. (B) Quantification of measurements of wild-type RBPMS-A (WT; maroon circles) and mutants (T/A; purple squares & T/E; cyan triangles) binding to RNA from (A) were fitted to a Hill equation. The apparent dissociation constants (K_d_) are listed below each gel (n = 3) and are for comparison between the mutants only. Error bars are SEM. (C) UV-crosslinking using recombinant wild-type (WT) RBPMS-A, T/A, T/E mutants. In vitro crosslinking experiments were carried out in the presence (+N.E.) or absence (-N.E.) of 20% HeLa nuclear extract in a serial dilution (1:4; 4 to 1 uM) as indicated by the black triangle with the µM amounts listed above. % cross-linked is normalized to 100% crosslinked at the highest concentration of WT (4 µM) ±SD.

### Phosphomimetic mutant of RBPMS-A interferes with higher-order oligomerization and RNA contact

Recently, we have found that RBPMS-A undergoes higher-order oligomerization that is dependent upon the last 20 amino acids of the C-terminal intrinsically disordered region (IDR) of RBPMS-A. Truncation of this region impaired RBPMS-A oligomerization, multivalent RNA-binding, and splicing activity *in vitro* (Yi Yang et al., manuscript in preparation). Therefore, we hypothesized that phosphorylation or phosphomimetic mutation of T113/118 might interfere with RNA-binding and activity by affecting oligomerization (and hence reducing the avidity of multivalent binding). An alternative, but not mutually exclusive, hypothesis is that the negatively charged phosphoT113/118 residues directly interfere with RNA-binding (via electrostatic effects) by the adjacent RRM. We decided to test both of these hypotheses. First, to investigate the potential effects upon multivalent RNA-binding, we used analytical ultracentrifugation (AUC) to analyze the sedimentation velocity (Figure 7A) and size distributions (Figure 7B) of wild-type rRBPMS-A and the rRBPMS-A T/A & T/E mutants. The WT and T/A mutant showed a mixture of dimers located at a sedimentation coefficient of ∼2.5S and higher-order oligomers located at a sedimentation coefficient of ∼18 (Figure 7B), with a preponderance of the larger assemblies. Remarkably, the RBPMS-A T/E mutant showed a drastic decrease in the proportion of oligomers (Figure 7B, cyan line) when compared to the wild-type (maroon line) and the T/A mutant (purple line). However, the dimerization capability of the T/E mutant appears to remain intact as indicated by SEC-MALS (Supplementary Figure S7A and B). These data suggest that the phosphorylated RBPMS-A shows reduced oligomerization which ultimately may lead to lower activity by reduced multivalent avidity for RNA, as indicated by EMSA and UV cross-linking.

Secondly, to explore whether phosphorylation at T113/118 might directly affect the ability of the RRM domain to interact with RNA, we used NMR spectroscopy to analyze truncated versions (amino acids 1-122) of rRBPMS-A WT and T/E mutant, which were double-labeled with ^13^C and ^15^N (Figure 7C, Supplementary Figure S6B). Chemical shift assignment was carried out using published data for a related protein as a reference (RBPMS2, BMBR Entry 19298, (22). The assignments were then extended to cover the disordered N-and C-terminal regions of RBPMS-A *ab initio*, facilitated by ^1^H/^13^C/^15^N experiments at a reduced temperature that gave optimal peak intensities for these mobile regions (see Methods). Following assignment, chemical shift differences (Δδ) between WT and T/E were quantified (Figure 7C) and linearly mapped from gray to red (greatest shift) onto a ribbon model of RBPMS-A bound to an RNA ligand (Figure 7D). The chemical shift differences were largest at or near the phosphomimetic mutations, as expected (Figure 7C). However, an inspection of the differences between the WT and T/E on the RRM domain itself revealed that a cluster of structurally contiguous residues in the RNA-binding site was strongly affected by the introduction of the phosphomimetic residues (Figure 7D). Further, the RNA-binding face is lined with basic residues that would be expected to preferentially bind acidic residues and/or phosphothreonine (Figure 7E). These findings further support our hypothesis that RBPMS-A’s RNA-binding region is directly affected by the addition of phosphomimetics (and by extension, phosphorylation) and that this effect likely reduces its RNA-binding affinity. Taken together, our data indicate that phosphorylation of RBPMS at T113/118 reduces RNA-binding by both directly interfering with the RNA-binding surface of the RRM domain and by reducing the avidity of multivalent RNA interaction due to the lower oligomerization state.

**Figure 7.**
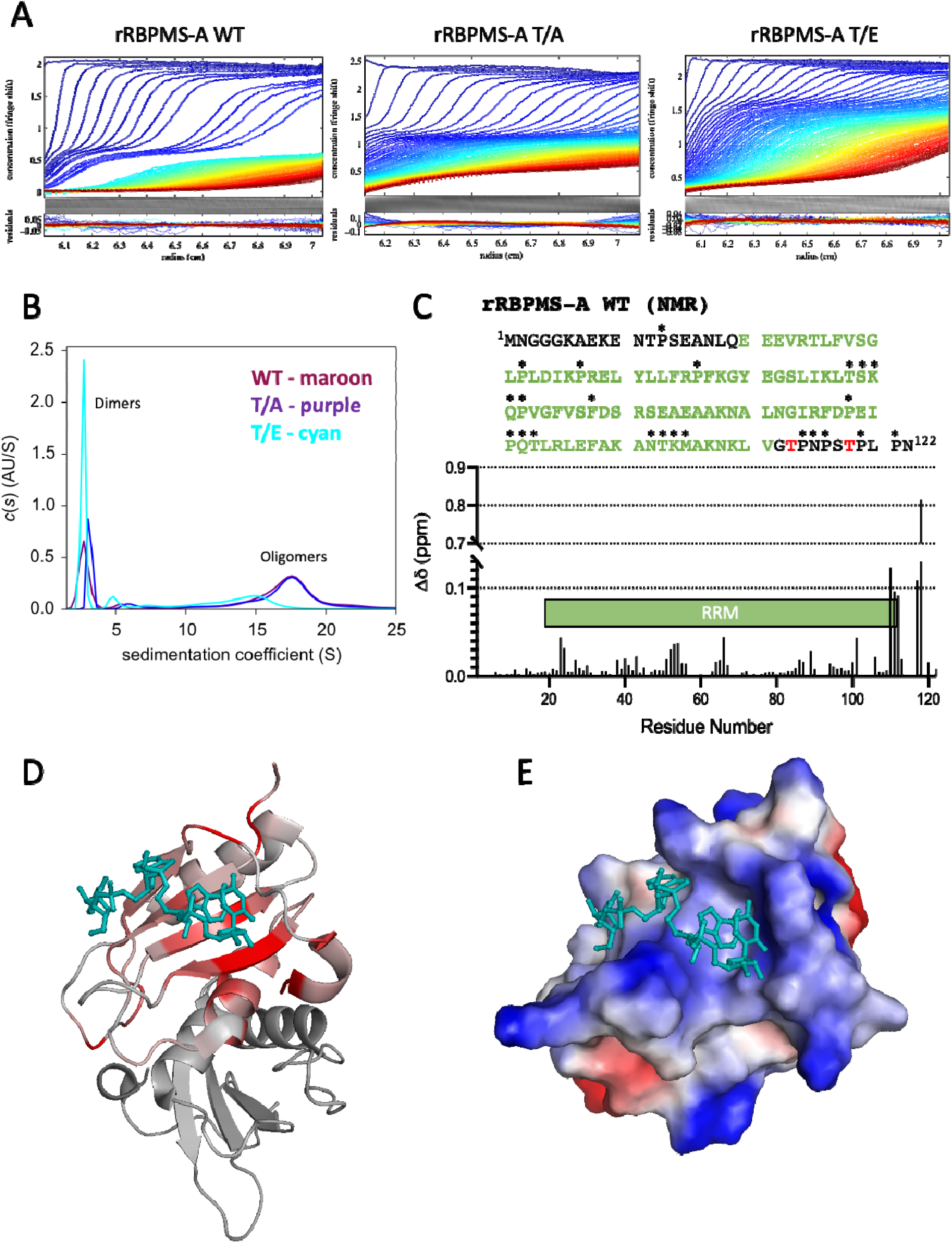
RBPMS-A Phosphomimetic Fails to Oligomerize and Directly Affects its RNA Binding Site. (A) Analytical Ultracentrifugation (AUC) sedimentation velocity acquired at 40,000 rpm for recombinant wild-type (WT) RBPMS-A and the T/A and T/E mutants at a concentration of 0.5 mg/mL. Radius (cm) is represented on the x-axis and the absorbance at 274 nm on the y-axis. Color temperature indicates the progress of time. (B) Size distribution profiles or continuous sedimentation coefficient distribution, c(S), curves of each recombinant protein determined from their sedimentation velocity in (A). WT=maroon line, T/A=purple line, T/E=cyan. The peaks around a sedimentation coefficient of 2 represent the protein in a dimeric form while the peaks around 18 represent an oligomeric form. (C) RBPMS-A WT truncated amino acid sequence used for NMR, mutations at T113 and T118 (shown in red) to glutamic acid were used for the T/E mutant for NMR. The RRM is shown in green. Unlabeled residues are marked with asterisks (*). Quantified chemical shift differences between RBPMS-A WT and T/E are shown in the graph. Cartoon of the RRM domain is annotated in green. (D) Chemical shift differences mapped onto an RBPMS-A dimer. Ribbon model of RBPMS is colored according to Δδ on a scale of 0-0.01, linearly mapped from gray to red. The darkest red residues are the ones that have shifted the most. Bound RNA is shown for illustrative purposes (cyan). (E) The electrostatic potential of an RBPMS-A monomer bound to RNA (cyan) in the same orientation as (D), blue representing basic residues and red representing acidic residues.

## DISCUSSION

Cell-specific master splicing regulators have been defined by the following criteria: 1) they are required for cell-type specification or maintenance, 2) their direct and indirect targets are functionally related and important for cell-type function, 3) they are likely to regulate the activity of other splicing regulators, 4) they have a wide dynamic range of activity, not limited by autoregulation, and 5) they are regulated externally from the splicing network via transcriptional control or post-translational modifications (PTMs) (44). Previous work in our lab identified RBPMS as a smooth muscle master splicing regulator by addressing criteria 1-4 (5). Regarding criterion 5, it was already known that RBPMS transcription is switched off during smooth muscle dedifferentiation (5). Here we provide evidence that the splicing regulatory activity of RBPMS protein is also directly down-regulated by phosphorylation, providing a rapid mechanism to switch off RBPMS activity as an early response during VSMC phenotypic modulation. A model summarizing our findings and their implications is shown in Figure 8. In differentiated contractile VSMCs (Figure 8, left) non-phosphorylated RBPMS exists as dimers, mediated by the RRM domain (43), which are further assembled into larger oligomers, mediated by the C-terminal IDR. RBPMS oligomers can bind multivalently to target RNAs with multiple CAC motifs, such as *Tpm1* exon 3, to promote splicing patterns characteristic of differentiated VSMCs. Upon vascular injury or some other perturbation, activation of signaling pathways converges upon ERK1 or 2, which then phosphorylate RBPMS at Thr113 and 118 (Figure 8, center). Phosphorylation of T113/118 shifts the equilibrium from higher order oligomers towards dimers, thereby reducing the avidity of multivalent RNA binding. In addition, the C-terminal region containing phospho-T113/118 acts as an RNA mimic, looping around and directly occluding the RNA binding surface of the RRM. Together, these two effects lead to reduced RNA binding by RBPMS and switching of its regulated AS events to the splicing pattern characteristic of proliferative VSMCs (Figure 8, right). While all of our data are consistent with this model, some elements of the model will require further experimental investigation (discussed below).

**Figure 8.**
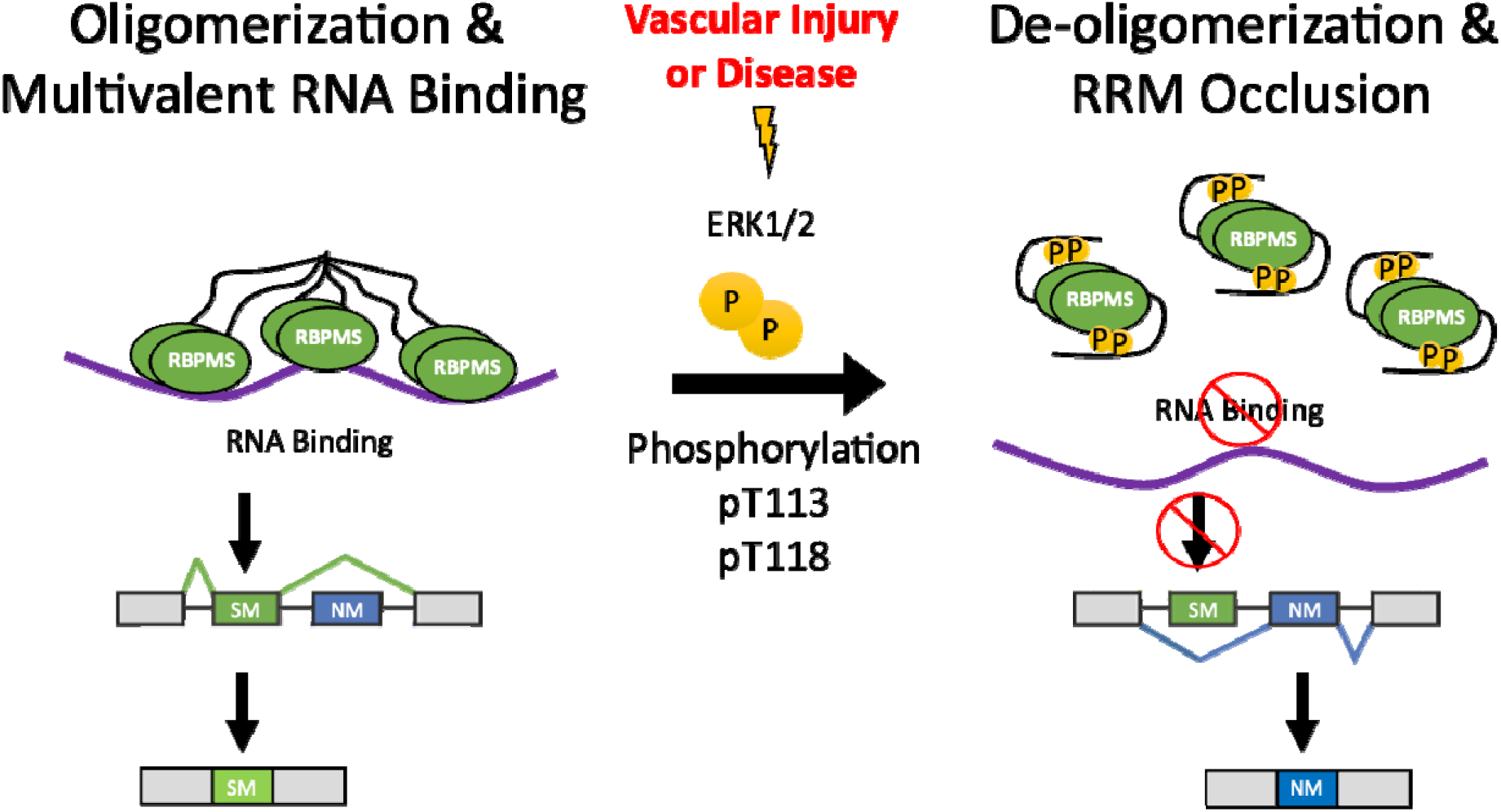
A Model of Splicing Inactivation of RBPMS-A via Phosphorylation. Vascular injury or disease activates ERK1/2 leading to the downstream phosphorylation of RBPMS-A at Thr113 and Thr118. Phosphorylation of RBPMS-A leads to de-oligomerization and RRM occlusion reducing RNA-binding and ultimately changing the splicing outcome, resulting in the increased inclusion of NM (non-smooth muscle) exons, found in the proliferative SMC cell type. In combination with the transcriptional downregulation of RBPMS, the rapid modulation of RBPMS via phosphorylation could play a vital role in phenotype plasticity during the vascular injury response and atherogenesis.

Multiple lines of evidence indicate that Thr113/118 are phosphorylated and that ERK2 (and possibly ERK1) is the responsible kinase (Figures 2-4, Supplementary Figure S1). However, our conclusions about the functional consequences of phosphorylation rest upon the use of phosphomimetic glutamate mutations at T113 and T118. Potential concerns about this approach are first that the mutation might impair the normal function of the non-phosphorylated threonines. The retention of full RBPMS activity upon deletion of aa112-120 (Supplementary Figure S1C) strongly argues against this possibility, suggesting instead that this region serves only as a platform for regulatory input by phosphorylation. A second concern is that Glu fails to completely mimic phospho-threonine. Glutamate residues have one negative charge at neutral pH, whereas phospho-threonine has between 1.5 and 2 negative charges. Phosphomimetic mutants are therefore likely to underestimate the consequences of phosphorylation, particularly where the phosphate group has a strong electrostatic effect. This is likely to be the case in the “RNA mimic" mode of action where the RNA binding surface of the RRM is occluded by the phosphopeptide (Figure 7C-E). While we do not yet understand the physical basis by which the phosphomimetics lead to de-oligomerization (Figure 7A and B), it seems likely that the T/E mutants underestimate the full consequences of phosphorylation, and that the down-regulation of RBPMS by phosphorylation might be more marked than we observe with phosphomimetic mutants. Consistent with this argument, we observed a complete loss of transfected RBPMS splicing activity upon co-transfection with MEK2-CA and ERK2 (data not shown), under conditions where RBPMS becomes fully phosphorylated (Figure 3E). However, the non-phosphorylatable T/A and T/A/S144A mutants were also inactive under these conditions, so the loss of activity could not be attributed to the phosphorylation of T113/118. A possible explanation is that ERK2 also phosphorylates and inactivates essential RBPMS coregulators such as RBFOX2 and ESRP2 which interact with RBPMS via its C-terminal IDR (Yang et al., manuscript in preparation) and are known to be ERK targets. In this case, we would need non-phospho mutants not only of RBPMS but also of all the essential co-regulators, to bypass the inhibitory effect of ERK2. As an alternative, future *in vitro* studies to analyze the effects of *bona fide* phosphorylation could use rRBPMS phosphorylated *in vitro*. However, a downside to this approach would be the heterogeneous and off-target phosphorylation that we observed with mass spectrometry (Figure 2D).

Phosphorylation of RNA binding proteins in general, and splicing factors in particular, have been widely documented. Phosphorylation can regulate RBP AS activity by affecting one or more of: RNA interactions; protein-protein interactions; subcellular localization; targeted degradation, reviewed in (45). In a limited number of cases, the consequences of phosphorylation are understood at a structural level (46). We observed some shift in localization of RBPMS T/E to the cytoplasm (Figure 5A, Supplementary Figure S5D), but the partial nature of this effect suggested that it was insufficient to have a major impact on nuclear RBPMS activity. Indeed, the loss of activity of RBPMS T/E in a cell-free splicing assay (Figure 5D) suggests that relocalization is unnecessary for the down-regulation of RBPMS splicing activity. Moreover, the reduced RNA binding of the phosphomimetic mutant (Figure 6) suggests that cytoplasmic RBPMS functions would also be down-regulated. This could be relevant for modulating the role of RBPMS as a translational regulator of lineage commitment in pluripotent cells by binding to 3’ UTRs of target RNAs (47). Notably, multiple RBPMS bands were observed in ES cells, which likely arose both from the known RBPMS isoforms, but also the phospho-variants described here.

We uncovered two potential mechanisms, which are not mutually exclusive, that lead to reduced RNA binding and loss of RBPMS splicing regulation: direct occlusion of the RNA binding surface of the RRM (Figure 7C-E), and reduced higher-order oligomerization (i.e., homomeric protein-protein interactions) in favor of dimers. While RRM occlusion is likely to affect all RNA binding events, reduced oligomerization is likely to disproportionately affect events involving multivalent RNA binding (such as *Tpm1* exon 3), reminiscent of the differential effects of TDP-43 oligomerization mutations upon binding to long-multivalent regions compared to short multivalent sites (48). We have recently found that RBPMS oligomerization, which can progress to liquid-liquid phase separation, is mediated by the C-terminal IDR (Yang et al., manuscript in preparation). A mutant with deletion of the C-terminal 20 amino acids unique to RBPMS-A lost splicing regulatory activity, oligomerization, interactions with other splicing co-regulators such as RBFOX2, ESRP2, RBM4, RBM14, RBM47, and binding to the multivalent *Tpm1* RNA substrate, similar to the phosphomimetic mutant (Figures 5-7). The phosphomimetic mutants showed differential loss of activity on different RNA substrates, generally having a greater impact on RBPMS repressed exons (Figure 1). It will be interesting in the future to investigate whether this is related to differential valency and distribution of binding sites around the different target exons, as observed for TDP43 mutants (48). An interesting possibility is that the two mechanisms that we have proposed for reduced RNA binding are actually physically coupled. In this scenario, the interaction of the phosphorylated T113/118 region with the RRM’s RNA binding face would directly impact the ability of the C-terminal tail to engage in homomeric interactions leading to oligomerization, as well as potentially heteromeric interactions with other proteins. To pursue this question, we will first need to develop a better understanding of the “molecular grammar” (49) underpinning RBPMS oligomerization and interactions with other proteins, and whether these hetero- and homo-typic interactions employ a common physical basis. A recent report showed that RNA binding and splicing activity of the splicing regulator SAM68 is also downregulated by phosphorylation in its N and C-terminal IDRs, flanking the canonical RNA binding domain (49a). However, in that case the phosphorylation occurs directly within Arginine-Glycine regions antagonizing additional RNA that are necessary for stable binding.

During VSMC phenotypic switching, adhesion receptors, extracellular matrix, and growth factors relay extracellular signal inputs (3). Of all the growth factors that activate the Ras/Raf/MEK/ERK pathway, platelet-derived growth factor (PDGF) is considered to be the master growth factor driving vascular smooth muscle cell (VSMC) dedifferentiation (50). PDGF is released at the site of tissue damage and stimulates the migration and proliferation of VSMCs (51–54). Additionally, PDGF-stimulated migration of VSMCs to the intima, the inner layer of cells lining the arterial lumen, is an initial step in atherosclerosis (3). Upon activation of this signaling pathway, differentiated VSMCs transition to a more motile and proliferative state. We propose that an early step in this transition is the inactivation of the smooth muscle cell master splicing regulator, RBPMS. Inactivation of RBPMS via phosphorylation by ERK1/2 would provide SMCs with a rapid and reversible mechanism to precisely regulate the activity of this protein. RBPMS expression is not detected in dedifferentiated VSMCs in either cell culture (5) or *in vivo* following a carotid ligation injury (55), where RBPMS is present in contractile but absent in proliferative SMCs (A. Jacob, & CWJS unpublished observations). Testing whether RBPMS phosphorylation occurs at an early stage of the injury response will require antibodies specific for T113/118 phosphorylated RBPMS. Finally, in light of the fact that the potent transcriptional regulator necessary for SMC phenotypic transition, KLF4, is directly regulated by PDGF through activation of ERK1/2 (56–59), it is an attractive hypothesis that this pathway also regulates the master splicing regulator of SMCs, RBPMS. The regulation of transcription and splicing in concert would ensure the rapid onset of transition from a differentiated to proliferative phenotype.

Together, the data presented here support a model for the rapid modulation of the master SMC splicing regulator, RBPMS, via phosphorylation in response to external signals during the vascular injury response and atherogenesis. This results in RBPMS de-oligomerization and RRM occlusion, thereby reducing RNA-binding and ultimately changing the splicing outcome, working in concert with transcriptional regulation, when smooth muscle cells transition to a more motile and proliferative phenotype from their previously quiescent state (Figure 8).

## DATA AVAILABILITY

The residues phosphorylated on RBPMS-A WT and T/A were identified via LC-MS/MS and mass spectrometry proteomics data have been deposited to the ProteomeXchange Consortium via the PRIDE (60) partner repository with the dataset identifier PXD034462.

## SUPPLEMENTARY DATA

Supplementary Data are available at NAR online.

## ACKNOWLEDGMENTS

The authors would like to thank Matthew Watson, Department of Biochemistry, University of Cambridge, for NMR protein expression protocols, and Remkes Scheele, Department of Biochemistry, University of Cambridge, for providing purified active rERK2, and expression vectors for MEK1-CA and ERK2-WT. We thank Alex Borodavka and Giselle Lee for helpful comments and feedback on the manuscript.

---

**Supplementary Figure S1.**
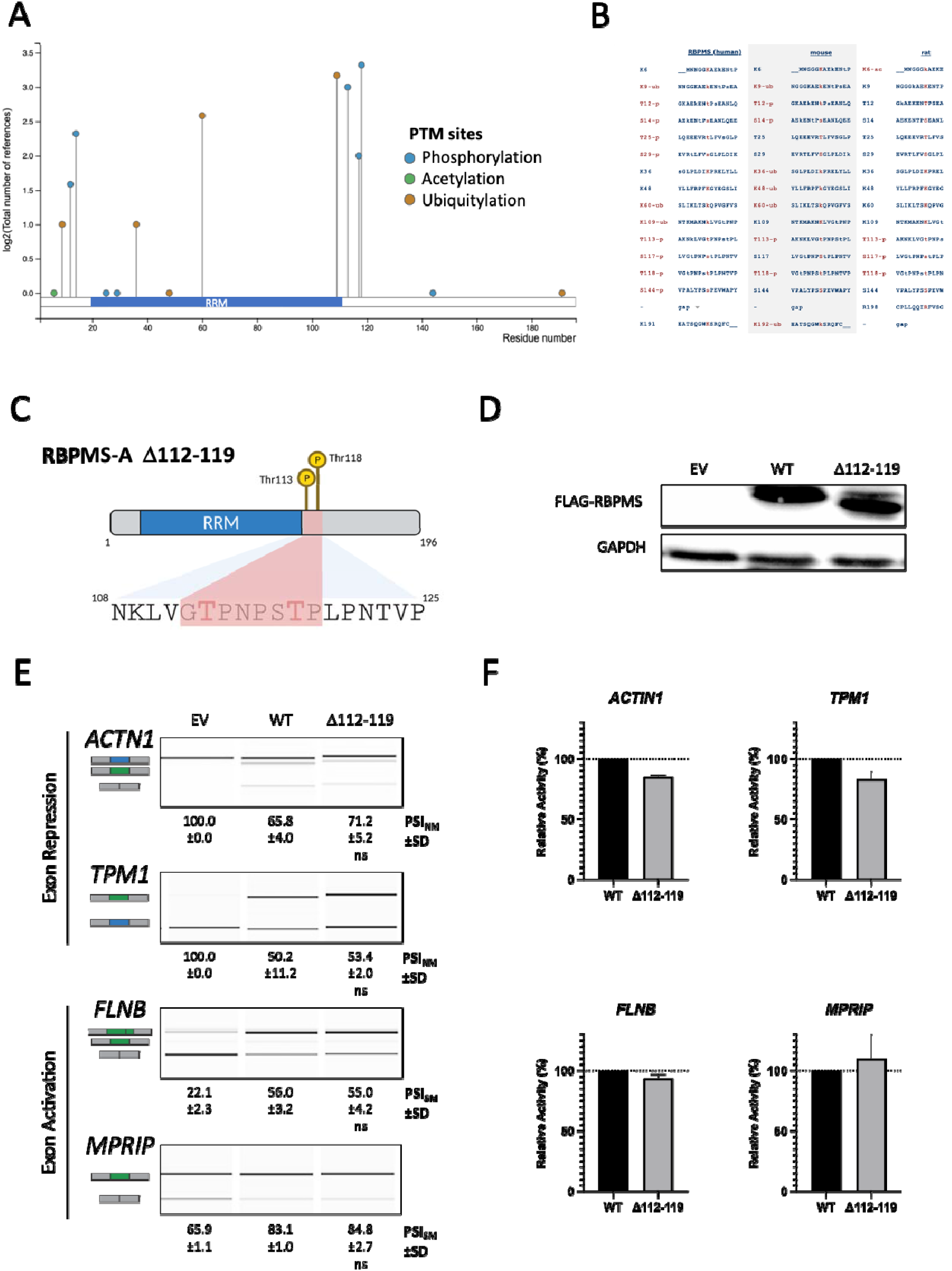
Post-translational Modifications Identified for RBPMS. (A) Post-translational modification location along RBPMS and the total number of references (published data sets), with modified RRM to reflect the correct length of the RRM. (B) Conservation of PTMs across mouse and rat species that have been observed thus far in published data sets. (A&B) Figures modified from (Hornbeck et al, 2015). https://www.phosphosite.org/proteinAction.action?id=2381994 (C) Graphical representation of RBPMS-A with the T113 and T118 highlighted in red. T/A is a double mutant of T113A and T118A, while T/E is a double mutant of T113E and T118E. Created with BioRender.com (D) Immunoblot of FLAG-RBPMS effectors used for splicing assays in (E). Whole-cell lysates were probed with antibodies to FLAG and GAPDH. GAPDH was used as a loading control. (E) Splicing analysis of the exon repression function of wild-type (WT) RBPMS-A and effectors in endogenous *ACTN1* and *Tpm1*, and its exon activation function in endogenous *FLNB* and *MPRIP* in HEK293T cells. RT-PCR followed by visualization on QIAxcel. Cartoons below the gene names represent the PCR products. The blue exon in the cartoon represents the NM (Non-smooth Muscle) exon, while the green exon represents the SM (Smooth Muscle) exon. PSI (Percent Spliced In) values are the mean ± SD (n = 3). A Student’s t-test was used to determine statistical significance between wild-type RBPMS-A and effector. EV=empty vector. (F) Bar graphs showing the percent relative splicing efficiency of each effector normalized to wild-type RBPMS-A (black bar) after background splicing was taken into consideration. Error bars indicate SD.

**Supplementary Figure S2.**
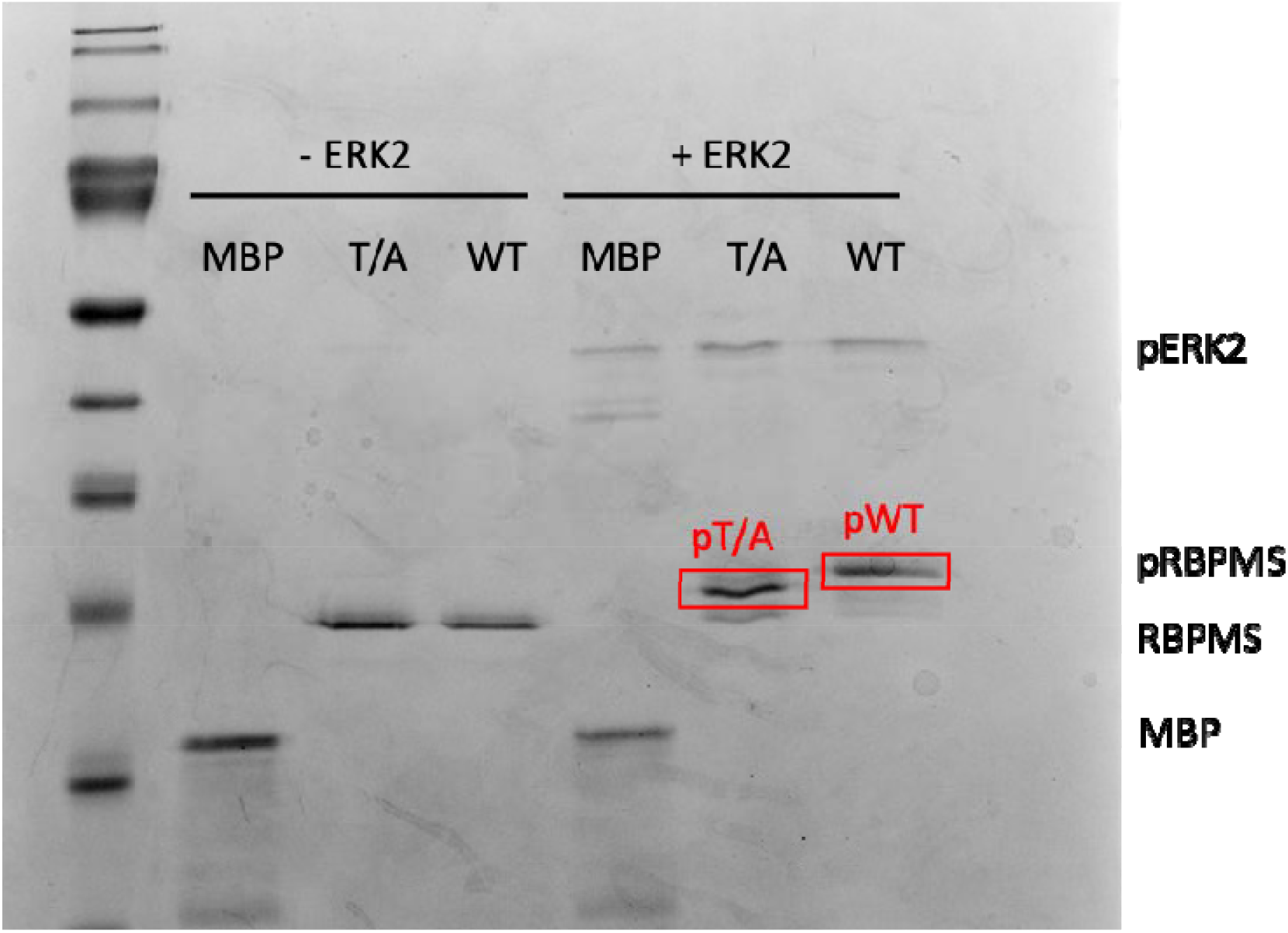
ERK2 Phosphorylates RBPMS-A. Recombinant wild-type (WT) RBPMS-A or RBPMS-A T/A (substrate; 1uM) were incubated with 200 nM highly active recombinant ERK2 in an *in vitro* kinase assay. Reaction was resolved on 20% SDS-PAGE and phosphorylated WT and T/A were excised from the gel (red rectangle) and sent for mass spectrometry. Myelin Basic Protein (MBP) was used as a positive control.

**Supplementary Figure S3.**
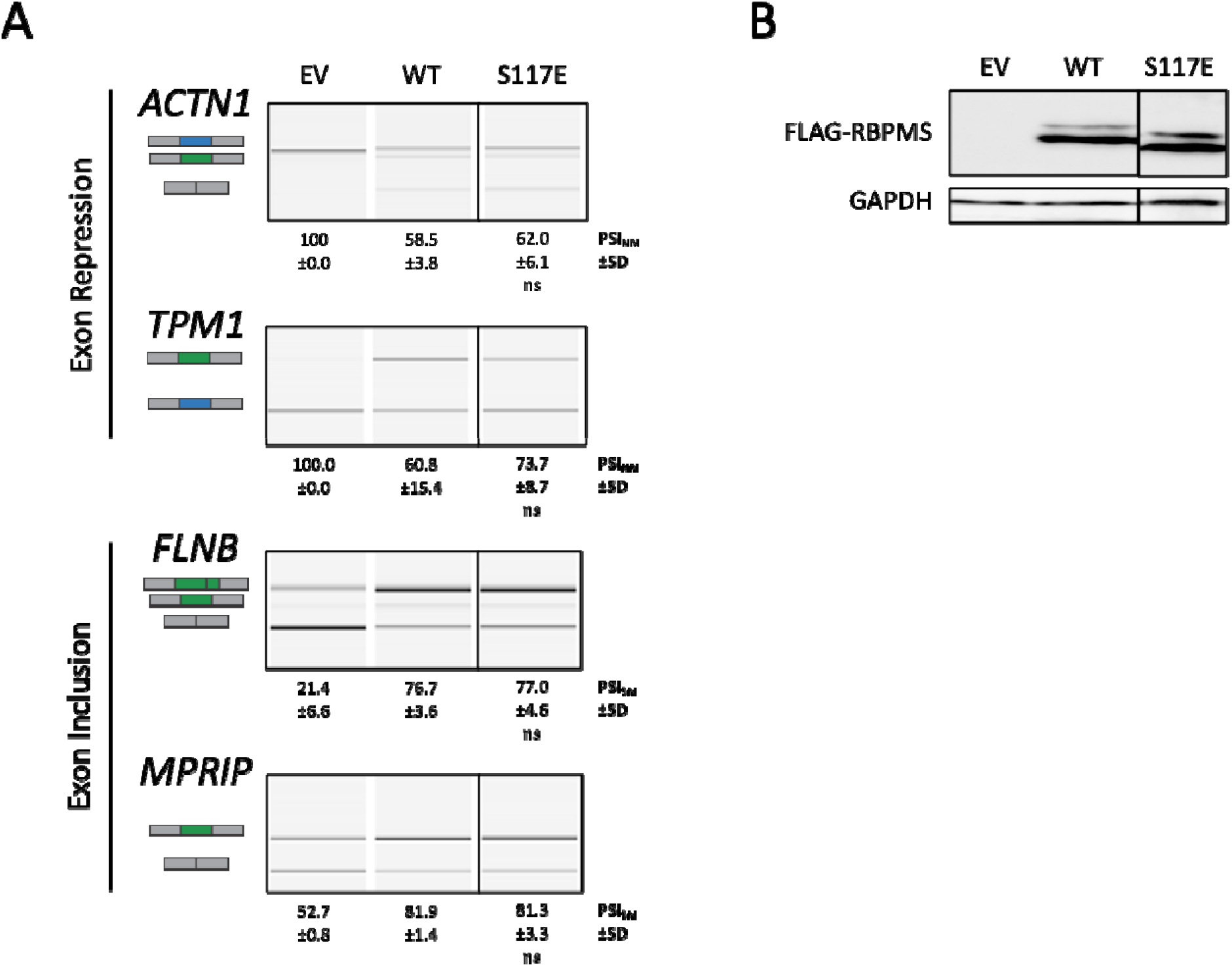
S117E Mutation has no Effect on RBPMS-A Splicing Activity. (A) Splicing analysis of the exon repression function of wild-type (WT) RBPMS-A and effectors in endogenous *ACTN1* and **Tpm1**, and its exon activation function in endogenous *FLNB* and *MPRIP* in HEK293T cells. RT-PCR followed by visualization on QIAxcel. Cartoons below the gene names represent the PCR products. The blue exon in the cartoon represents the NM (Non-smooth Muscle) exon, while the green exon represents the SM (Smooth Muscle) exon. PSI (Percent Spliced In) values are the mean ± SD (n = 3). A Student’s t-test was used to determine statistical significance between wild-type RBPMS-A and effector. EV=empty vector (B) Immunoblot of FLAG-RBPMS effectors used for splicing assays in (B). Whole-cell lysates were probed with antibodies to FLAG and GAPDH. GAPDH was used as a loading control.

**Supplementary Figure S4.**
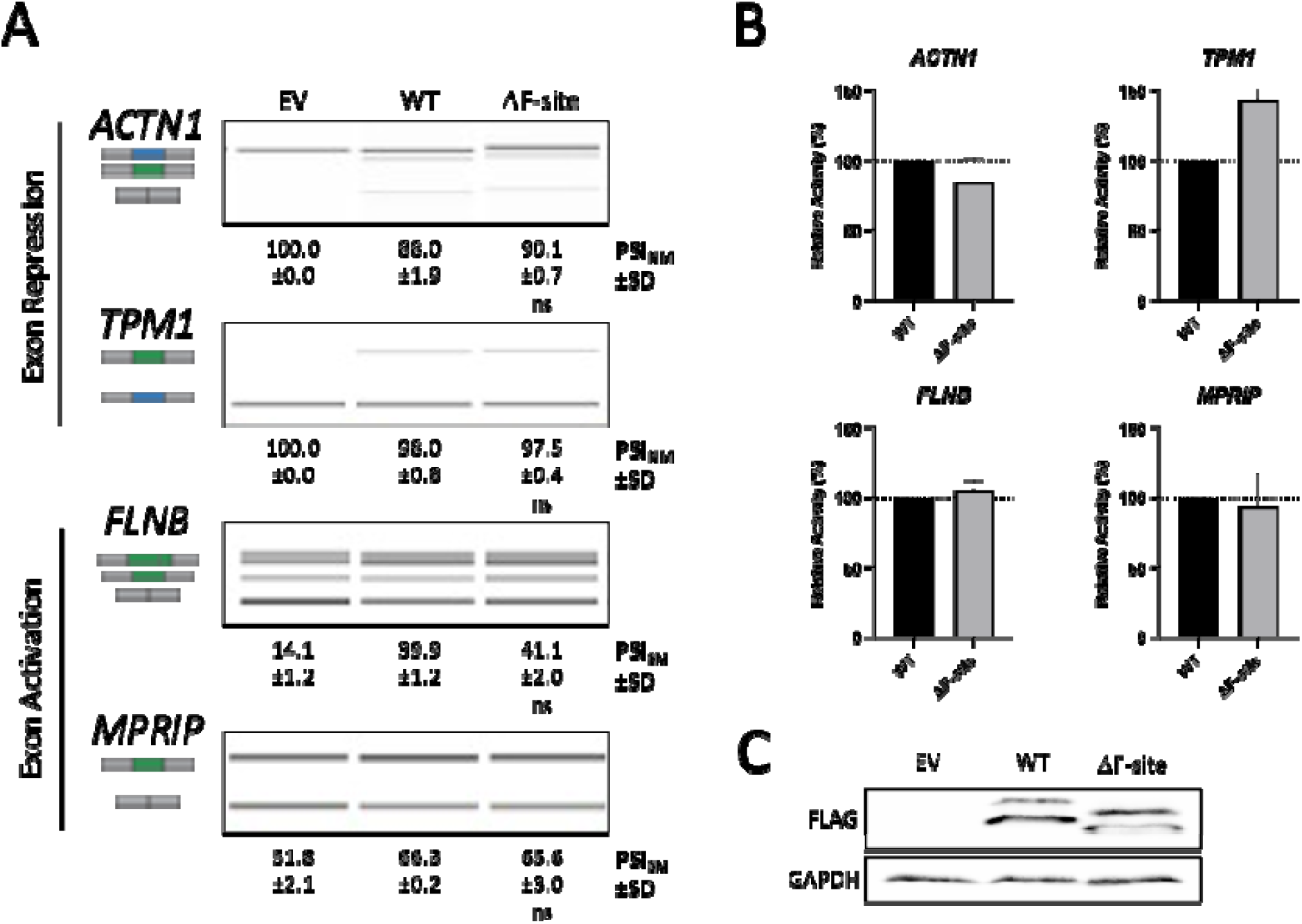
Deletion of the ERK2 Docking Site (F-site) has no Effect on Splicing Activity. (A) Splicing analysis of the exon repression function of wild-type (WT) RBPMS-A and effectors in endogenous *ACTN1* and **Tpm1**, and its exon activation function in endogenous *FLNB* and *MPRIP* in HEK293T cells. RT-PCR followed by visualization on QIAxcel. Cartoons below the gene names represent the PCR products. The blue exon in the cartoon represents the NM (Non-smooth Muscle) exon, while the green exon represents the SM (Smooth Muscle) exon. PSI (Percent Spliced In) values are the mean ± SD (n = 3). A Student’s t-test was used to determine statistical significance between wild-type RBPMS-A and effector. EV=empty vector. (B) Bar graphs showing the percent relative splicing efficiency of each effector normalized to wild-type RBPMS-A (black bar) after background splicing was taken into consideration. Error bars indicate SD. (C) Immunoblot of FLAG-RBPMS effectors used for splicing assays in (C). Whole-cell lysates were probed with antibodies to FLAG and GAPDH. GAPDH was used as a loading control.

**Supplementary Figure S5.**
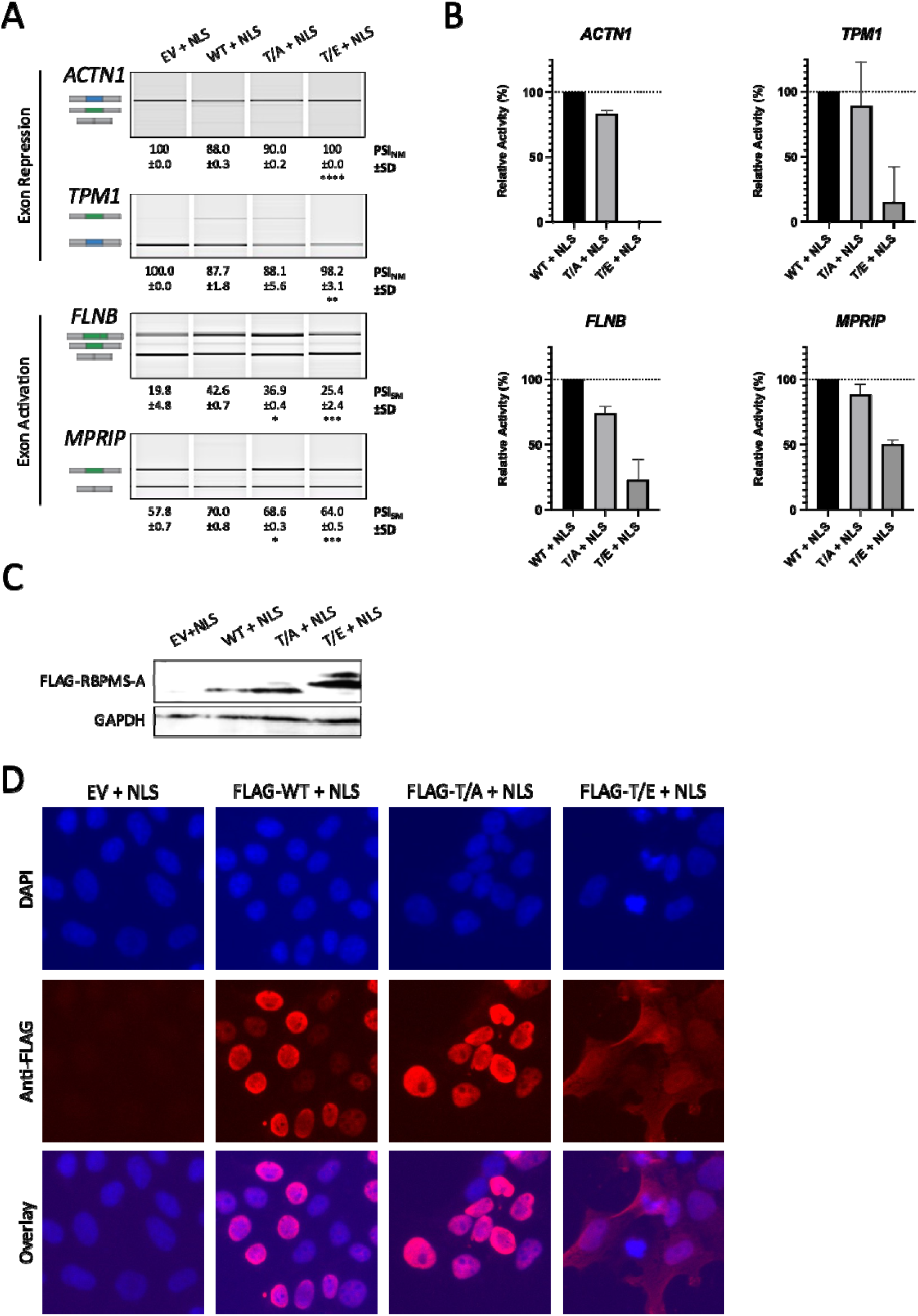
Exogenous NLS Fails to Rescue the Splicing Activity of RBPMS-A T/E. (A) Splicing analysis of the exon repression function of wild-type (WT) RBPMS-A and effectors in endogenous *ACTN1* and *Tpm1*, and its exon activation function in endogenous *FLNB* and *MPRIP* in HEK293T cells. RT-PCR followed by visualization on QIAxcel. Cartoons below the gene names represent the PCR products. The blue exon in the cartoon represents the NM (Non-smooth Muscle) exon, while the green exon represents the SM (Smooth Muscle) exon. PSI (Percent Spliced In) values are the mean ± SD (n = 3). A Student’s t-Test was used to determine statistical significance between wild-type RBPMS-A and effector, indicated by *p < 0.05, **p < 0.01, ***p < 0.001, ****p < 0.0001. (B) Bar graphs showing the percent relative splicing efficiency of each effector normalized to wild-type RBPMS-A (black bar), after background splicing was taken into consideration. Error bars indicate SD. (C) Immunoblot of FLAG-RBPMS effectors used for splicing assays in (A). Whole-cell lysates were probed with antibodies to FLAG and GAPDH. GAPDH was used as a loading control. (D) Immunofluorescence microscopy images of HEK293T cells overexpressing wild-type (WT) RBPMS-A and the T/A and T/E mutants all containing an exogenous nuclear localization signal (NLS). All images were taken with the same parameters using a 40X objective. Top row shows DAPI staining for DNA, middle row shows staining for FLAG-tagged effectors, and bottom row is an overlay of the DAPI and FLAG images.

**Supplementary Figure S6.**
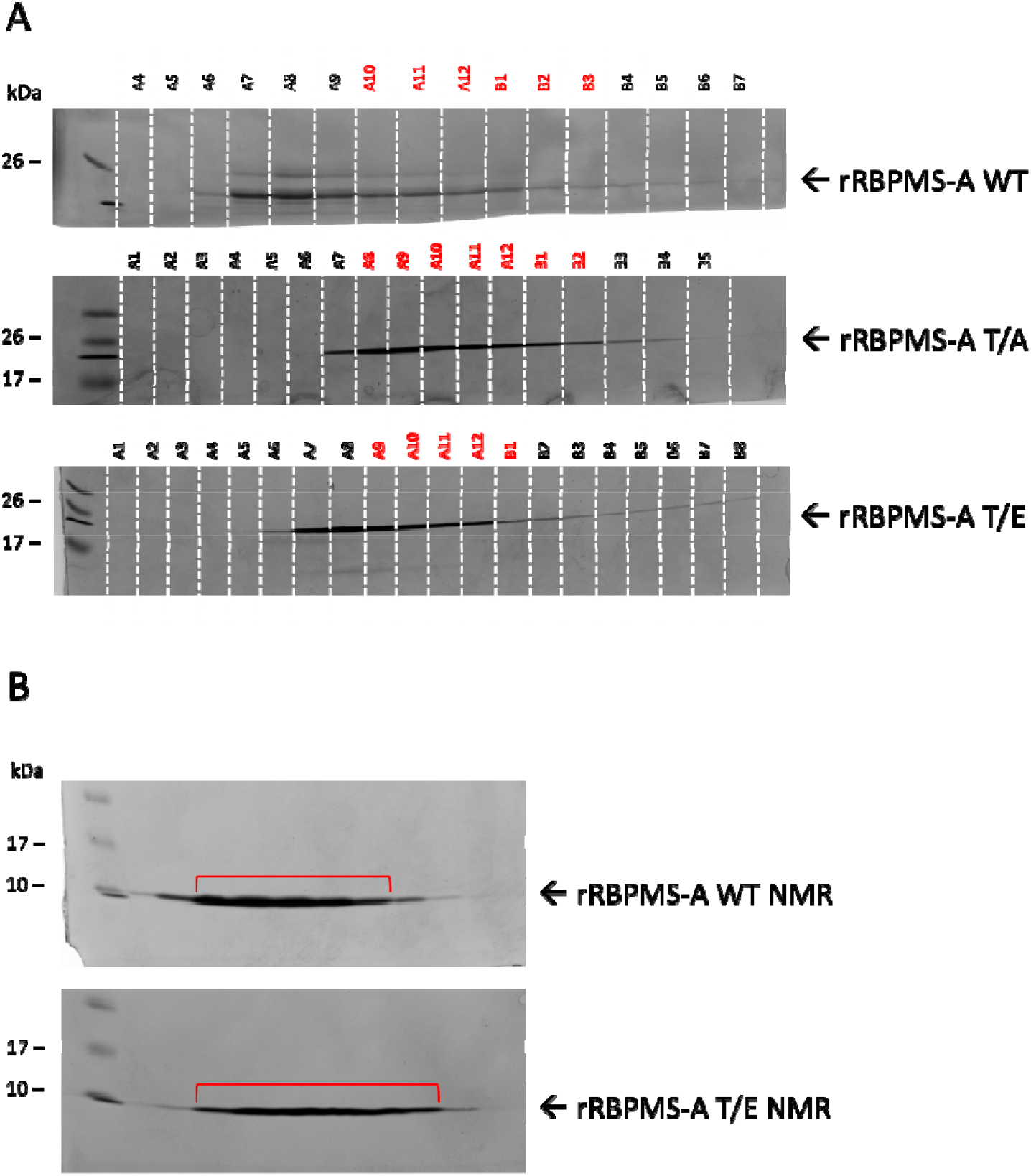
Purification of Recombinant RBPMS-A. (A) Final Mono Q^TM^ anion exchange chromatography purification of recombinant RBPMS wild-type (WT), T/A, and T/E mutants following TEV protease treatment to remove His-tag. See methods for a detailed purification method. Fractions highlighted in red for each protein were combined and used for follow-on experiments. (B) Final HiTrap^TM^ Heparin chromatography purification of recombinant double-labeled (^13^C/^15^N) RBPMS-A aa1-122 WT and T/E mutant. See methods for detailed labeling and purification method. Fractions highlighted by the red bracket were combined and used for follow-on NMR experiments.

**Supplementary Figure S7.**
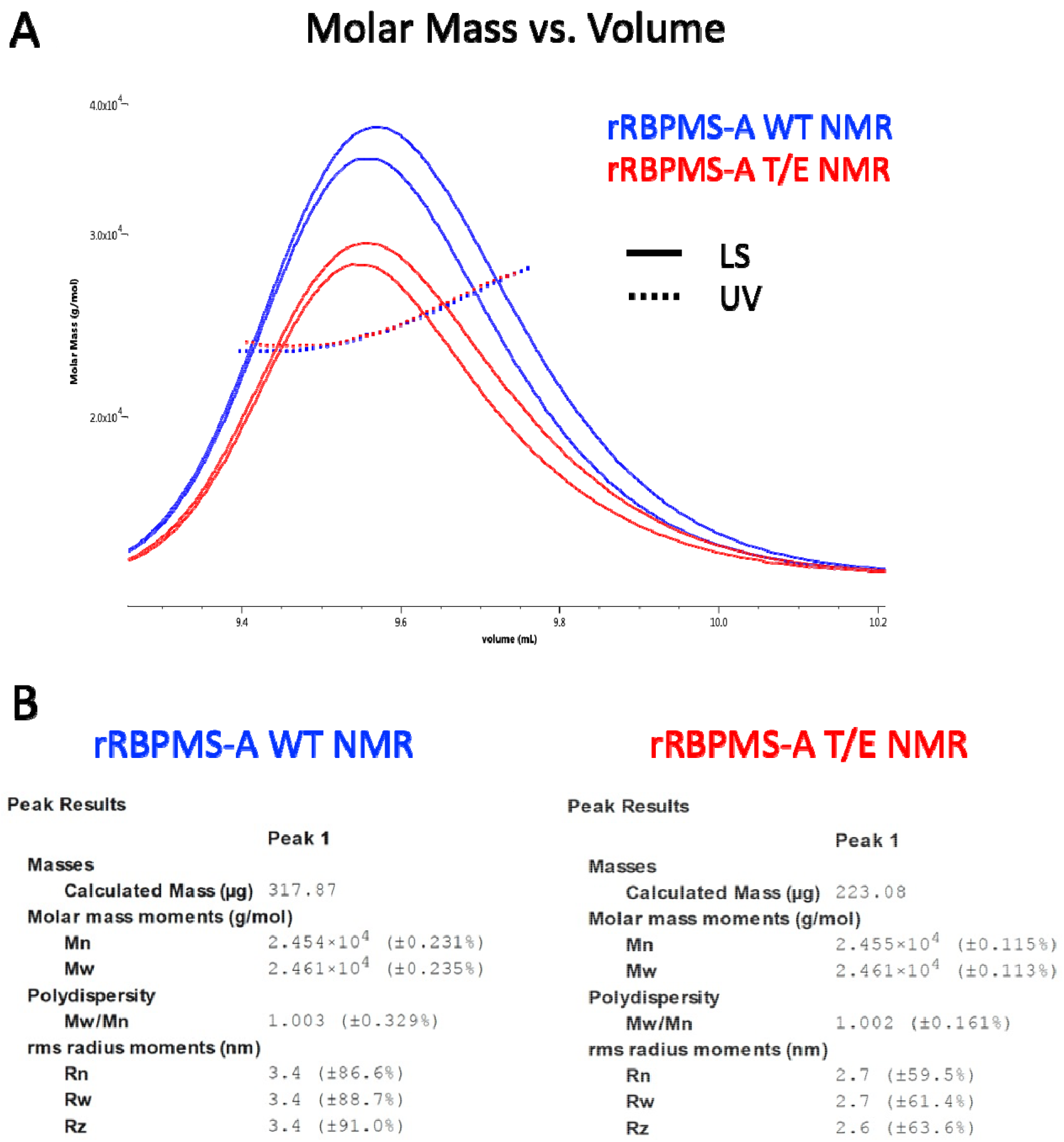
Truncated rRBPMS-A Proteins for NMR Remain Dimeric. Size exclusion chromatography multi-angle light scattering (SEC-MALS) of truncated NMR proteins. Conditions for both samples: 10 mM HEPES pH6.8 50 mM KCl. LS = light scattering; UV = ultraviolet absorbance (A) Chromatogram traces of each protein (B) MALS data output for each of the peaks in (A).

